# Brain-enriched coding and long non-coding RNA genes are overrepresented in recurrent autism spectrum disorder CNVs

**DOI:** 10.1101/539817

**Authors:** Hamid Alinejad-Rokny, Julian I.T. Heng, Alistair R.R. Forrest

## Abstract

Autism spectrum disorder (ASD) is a neurodevelopmental disorder with substantial phenotypic and etiological heterogeneity. It is estimated that 10-20% of cases are due to copy number variations (CNVs). Here we apply a newly developed CNV association tool (SNATCNV) to reanalyse CNV data from 19,663 autistic and 6,479 control subjects from the AutDB database. We demonstrate that SNATCNV outperforms existing CNV association methods by finding smaller genomic regions that better discriminate cases and controls. By integrating data from the FANTOM5 expression atlas we show that both known ASD causal genes identified by the SFARI and MSSNG consortia and genes within the CNVs identified by SNATCNV have brain enriched expression patterns; both brain-enriched coding and long-non-coding RNA genes are over-represented. We provide full lists of these brain enriched coding and lncRNA genes as a resource to the research community. We also go on to show that each CNV region is associated with a distinct set of phenotypes, that some are sex biased and highlight one deleted region where a brain-enriched lncRNA is the only gene present. Our analyses identify 47 high confidence ASD associated CNV regions and identifies brain-enriched genes which underlie this neurodevelopmental disorder.

## Introduction

Autism spectrum disorder (ASD) is collectively used to describe a group of neurodevelopmental disorders with a spectrum of phenotypes including impaired social interaction, repetitive behaviours, learning and speech impediments. There is a strong genetic component to ASD with a recent study of familial risk estimating heritability at 83%, and suggesting the majority of ASD risk is attributable to genetic factors (1, 2). The types of genetic lesions that lead to ASD are heterogeneous, and range from highly penetrant single gene mutations and copy number variations (CNV), through to weakly penetrant risk alleles identified by GWAS (3). Of note, recent studies reported by the SFARI (4) and MSSNG (5) consortia have identified 98 recurrently mutated, highly penetrant ASD candidate coding genes. Further, an estimated 15-25% of cases are attributable to *de novo* mutations (3), the majority of which correspond to CNVs.

There are at least 14 independent CNVs that have been previously associated with ASD (6, 7). Several are associated with distinct phenotypes, for example, the Prader-Willi/Angelman syndrome region on 15q11-q13 (8) is associated with obesity and ASD, while 1q21.1 is associated with intellectual disability (ID) and ASD; terminal deletions of 2q and 22q (9); and microdeletion and microduplication of 16p11.2 are associated with developmental delay (DD), ID and ASD (10). Despite the importance of CNVs to ASD causation, there remains a need to clarify what fraction of ASD is actually due to CNVs as estimates range greatly, from 10-20% (11–13). Additionally to understand the causative genes within these CNV regions, improved methods are needed to focus on the critical regions most strongly associated with ASD and prioritise the (often hundreds of) genes contained within each.

Here we have applied a novel CNV association tool, SNATCNV (Single Nucleotide Association Test for CNVs) to a combined CNV dataset curated by the Simons Foundation Autism Research Initiative (SFARI) containing 19,663 autistic and 6,479 control samples from multiple published data sets (14). Benchmarking against PLINK (15) showed that SNATCNV was able to identify smaller critical regions that better discriminate ASD cases from controls. Using the FANTOM5 (Functional Annotation of the Mammalian Genome 5) (16) CAGE associated transcriptome atlas we then show that critical regions identified by SNATCNV contain substantially more brain-enriched coding and long-non-coding RNA genes than expected based on the genome-wide occurrence of these genes. Notably, such gene enrichment was not observed for regions identified by PLINK. Furthermore, we attribute distinct phenotypes and ontological traits to distinct ASD CNVs. To our knowledge, our results represent the first account of recurrent ASD associated CNV regions containing a significant overrepresentation of genes with nervous system enriched expression profiles.

We present a summary of the regions identified in our analysis, assessment of their novelty, candidate ASD associated coding and long-non-coding RNA genes within the regions, phenotypes associated with specific CNVs and validation of the identified regions in two independent cohorts of ASD patients. Our results show that these regions are enriched for brain-specific genes, and that specific gene ontology terms and phenotypes are associated with different regions. This highlights the spectrum of genetically diverse etiologies currently grouped together as ASD.

## Results

### Robust identification of ASD associated CNVs

To robustly identify recurrently deleted and duplicated regions significantly associated with ASD, we developed a novel tool called SNATCNV (Single Nucleotide Association Test CNV) and applied it to a cohort of 19,663 autistic and 6,479 non-autistic individuals (http://autism.mindspec.org/autdb; download date: July 2016). For every position in the genome, SNATCNV counts how often the nucleotide is either deleted or duplicated in the case and control CNVs. It then uses the one-tailed Fisher’s exact test (**Supplementary figure S1**) to determine whether the base is significantly more frequently duplicated or deleted in cases or controls. As an example of our output, **Figure 1a** and **b** shows the distribution of *P* values for deletions and duplications in ASD found on chromosome 17, respectively. To identify significant regions, we calculated *P* values for 500,000 random permutations of case/control labels to estimate the probability that an association emerges by chance (see **Methods** for further details). To filter for significant associations, users can specify a confidence interval based on random permutations (the dashed lines in **Figure 1a** and **b** show regions identified at the 95% and 99.9999% confidence intervals; also see **Methods** and **Supplementary figures S2 and S3, Supplementary table S1**).

**Figure 1.**
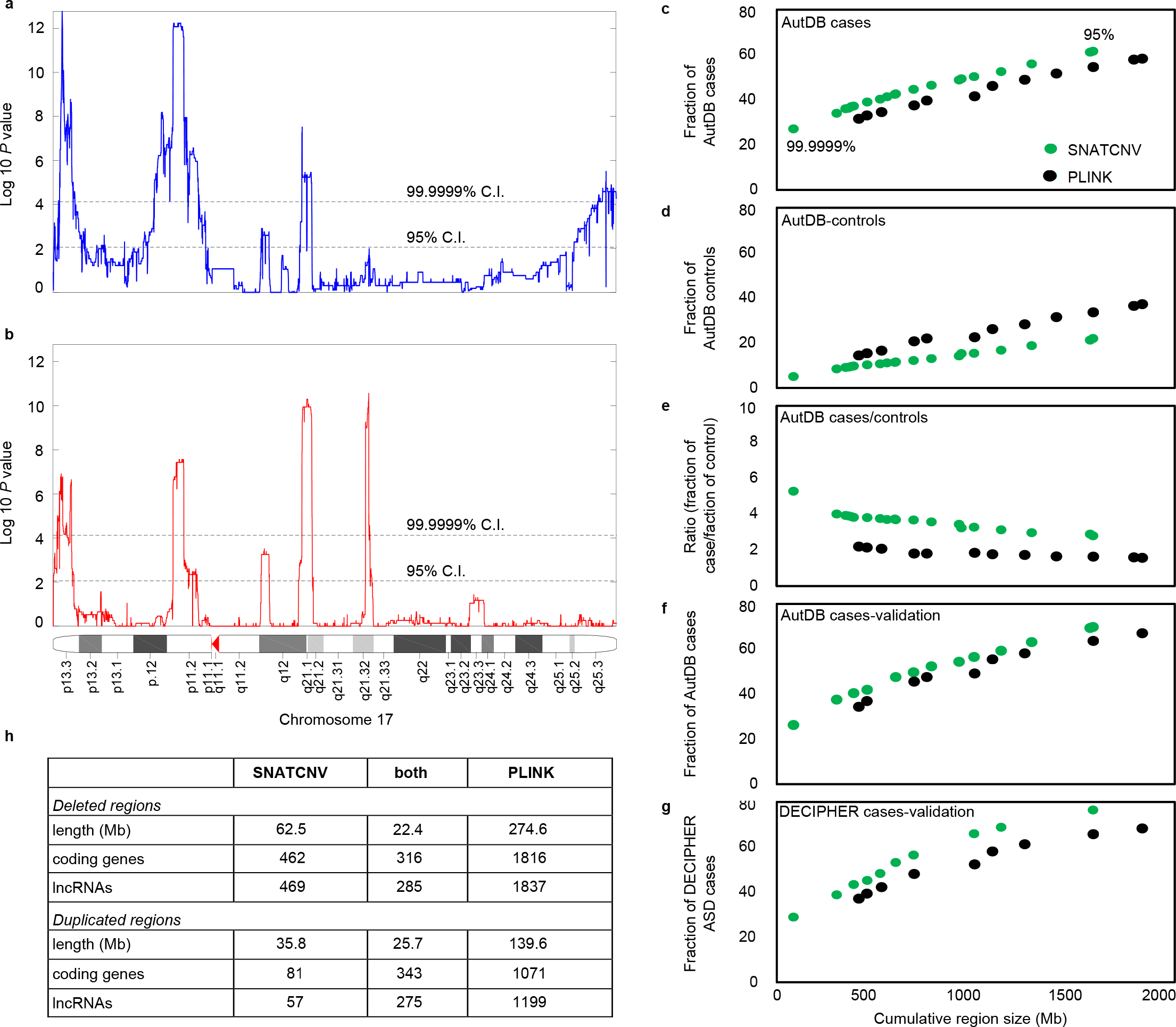
Identification of ASD associated regions using SNATCNV. Distributions of log10 *P* values found in ASD by SNATCNV comparing **a)** duplications and **b)** deletions on chromosome 17. Black dashed lines indicate our threshold for significant regions at two confidence intervals. X-axis indicates genomic loci and y-axis indicates logarithm 10 *P* value. *P* value calculated through Fisher’s exact test. Find more figures in **Supplementary figure S3**. Right panels (**c-g**) compare the signals obtained by SNATCNV (green) and PLINK (blank). **c)** Plot shows the fraction of case samples covered by SNATCNV’s or PLINK’s regions as a function of combined region size. **d)** As in c, except showing the fraction of controls covered. **e)** Shows the ratio of fraction of ASD samples vs fraction of control samples from c and d. Validation of the regions identified above using an independent cohort of **f)**1441 ASD individuals from the 2018 AutDB update and **g)**549 ASD individuals from DECIPHER.

To evaluate the performance of SNATCNV, we compared the regions identified across a range of confidence intervals to regions identified by the most commonly used CNV tool PLINK (15). Shown in **Figure 1c-e**, SNATCNV identified smaller regions that account for a significantly larger proportion of cases and significantly fewer controls than those identified by PLINK at all tested confidence intervals. At the most stringent threshold of 99.9999% C.I., 27% of ASD cases and 5.1% of controls overlap the identified regions thus the regions identified by SNATCNV are 5.3 fold more likely to be found in cases than controls. This is in comparison to only 2.2 fold for PLINK at its most stringent threshold.

Importantly, the regions identified in this cohort by SNATCNV explain a greater proportion of ASD case CNVs from two additional independent cohorts than those found by PLINK (**Figure 1f** - 1,441 additional ASD cases in the newest release of AutDB (6 Jun 2018) and **Figure 1g** - 549 ASD cases contained within DECIPHER (Database of Chromosomal Imbalance and Phenotype in Humans Using ENSEMBL Resources; 1 Feb 2017) (17)).

We also compared SNATCNV performance against the next most commonly used CNV association tool ParseCNV (18). ParseCNV requires the input data be in PennCNV format therefore we used a CNV dataset provided on the ParseCNV website (http://parsecnv.sourceforge.net) in PennCNV format that includes 785 ASD cases and 1,110 control individuals. Here again, SNATCNV outperformed both PLINK and ParseCNV (**Supplementary figure S4**). For a detailed comparison of features and limitations of these and other CNV tools see **Supplementary table S2**.

### Overrepresentation of brain-enriched genes in Autism associated CNV regions

Next, we focussed on 118 CNV regions identified by SNATCNV at the highest threshold of 99.9999% C.I., corresponding to 56 deleted and 62 duplicated regions, for further analysis of gene content. We used the FANTOM5 CAGE associated transcriptome (19) to identify 1,025 coding and 926 lncRNA genes contained within the 118 regions and examined their expression patterns across 347 sample types (**Figure 2a**). Hierarchical clustering of the genes based on their expression profiles revealed a substantial proportion of genes were highly expressed in the brain while smaller clusters of genes were highly expressed in blood, T-cells and tissues of the reproductive system.

**Figure 2.**
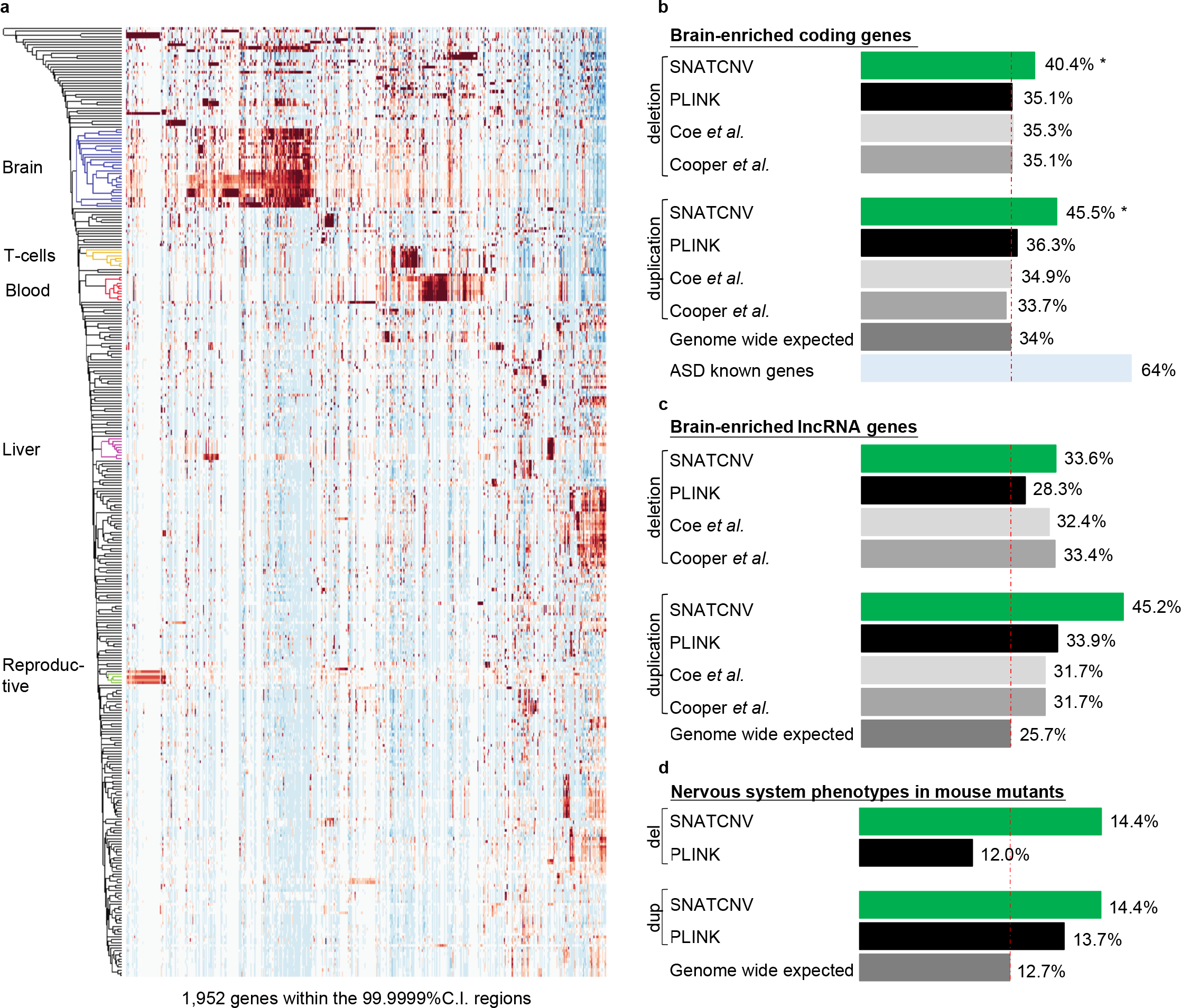
ASD associated CNVs identified by SNATCNV contain proportionally more nervous system enriched genes than expected. **a)** Hierarchical clustering of 1,952 coding and lncRNA genes from the highly significant regions based on their average expression across 347 sample ontologies (using the FANTOM CAT expression atlas (19)). Intensity represents the average level of gene expression across all samples annotated with the same sample ontology. **b)** Fraction of coding genes with brain enrichment found in the recurrently deleted and duplicated regions identified in the AutDB cohort using SNATCNV and PLINK at their highest thresholds. The dashed red line indicates the fraction of all genes in the genome with brain-enriched expression **c)** as in b, except for brain-enriched long non-coding RNA genes. **d)** as in b except plotting the fraction of mouse protein coding homologs in the regions that are associated with nervous system phenotypes in MGI.

To test whether the ASD associated CNV regions contained an overrepresentation of genes expressed in distinct tissues we next calculated the fraction of all coding and lncRNA genes in the genome with enriched expression in each tissue and compared the fractions to that observed within the CNV regions (see **Methods** and **Supplementary table S3**). Through this analysis, only brain-enriched genes were significantly overrepresented within ASD CNVs (**Figure 2b, c and Supplementary table S3**). We note, this enrichment was also observed when considering different sub structures of the brain. (**Supplementary table S4**).

Supporting the notion that brain-enriched genes are more likely to be causative in ASD we note that 64% of the ASD genes identified by SFARI and MSSNG are brain-enriched by our criteria, this is well above the genome wide expectation of 35% (**Figure 2b**). In contrast the regions identified by PLINK and the two largest studies of ASD, intellectual disability and developmental delay to date, Coe *et al.* and Cooper *et al.* (20, 21), showed no such brain enrichment. Although enrichment of brain enriched genes in deleted regions has been observed by Pinto *et al*. (22), to our knowledge this is the first report of overrepresentation of brain-enriched genes in recurrently deleted and duplicated CNV regions associated with ASD. Further supporting our analyses, the CNV regions identified by SNATCNV contain an overrepresentation of homologs of coding genes that cause nervous system phenotypes when mutated in the mouse (http://www.informatics.jax.org/mp/annotations/MP:0003631, 12 June 2018) (**Fig. 2c**).

### Brain-enriched coding genes within highly significant regions are associated with ASD specific biological processes and phenotypic ontologies

We next used WebGestalt (23) to investigate whether the 346 brain-enriched coding genes from the deleted and duplicated regions were associated with specific gene ontology (GO), human phenotype ontology (HPO) or disease terms (DisGeNet and GLAD4U) (24–26) (**Supplementary table S5;** all brain-enriched coding genes were used as a background). As expected, candidate genes in the CNV regions were more likely to be annotated with the ASD related disease ontology terms ‘autism spectrum disorder’, ‘intellectual disability’, ‘speech disorders’, ‘Angelman syndrome’, ‘Prader-Willi syndrome’ and ‘Rett syndrome’. Similarly overrepresented human phenotype ontology terms included 'Autism’ and ‘Neurological speech impairment’. Interestingly the most common overrepresented gene ontology term for 19 genes was ‘axon development’, which is a key developmental process in the etiology of ASD (27). Together this supports the biological relevance of the CNV regions identified by SNATCNV.

### Independent regions and dependent regions

Manual examination revealed some of the 118 regions were closely separated on the genome and were likely larger regions that were split by SNATCNV. This appears to be due to small regions of common CNV found in both the controls and cases which cause the *P* value to lose significance (see **Supplementary figure S5** as an example). Given this observation we next examined the independence of each of the 118 regions.

For each region we removed all case and control samples that overlapped the region and then recalculated the *P* value for the remaining 117 regions. This was repeated for all 118 regions (see **Supplementary figures S6** and **7** for *P* values after removal of deleted and duplicated regions, respectively). For 36 regions, removal of cases and controls found in any other region still yielded a significant *P* value (< 0.002), these we refer to as independent regions. For the remaining regions, 51 were removed as they lost significance when the cases and controls from the 36 independent regions were withheld. A further 31 corresponded to co-dependent scenarios where removal of all cases and controls from one region or another led to reciprocal lost significance. These 31 co-dependent regions were then merged into 11 superset independent regions (see **Supplementary figure S8, Supplementary table S6**). Thus we focus on 47 independent regions, comprising 36 non-merged independent regions and 11 independent merged superset regions (formed from 31 co-dependent regions) (**Table 1**). Note repeating the heatmap from **Fig. 2a** using the 1,649 genes contained within the 47 regions confirmed they are similarly enriched for brain genes (**Supplementary figure S9**).

**Table 1.** Brain enriched genes in the 47 recurrent CNV regions identified by SNATCNV analysis of the AutDB dataset (also see **Supplementary table S9**).

| Locus | chromosomal coordinates (hg19) | type | #case | #control | P value | #Brain-enriched / total |  | #Mice neuro phenotype |
| --- | --- | --- | --- | --- | --- | --- | --- | --- |
|  |  |  |  |  |  | coding | lncRNA |  |
| 1p36.32 | chr1:10001-10270615 | Del | 240 | 41 | 5.1E-06 | 52/155 | 42/152 | 16 |
| 1q21.1 | chr1:146113643-147727385 | Dup | 192 | 12 | 2.6E-17 | 4/10 | 6/15 | 0 |
| 1q21.1 | chr1:146113643-147870095 | Del | 244 | 14 | 6.2E-18 | 3/12 | 4/23 | 0 |
| 2q23.1 | chr2:148669357-148995585 | Del | 50 | 1 | 6.7E-06 | 3/3 | 2/2 | 2 |
| 2q37.3 | chr2:242930600-243007359 | Del | 63 | 1 | 1.7E-07 | 0 | 1/3 | 0 |
| 2p16.3 | chr2:51105853-51359022 | Del | 160 | 12 | 2.2E-10 | 1/1 | 1/1 | 1 |
| 3q29 | chr3:195745603-197355603 | Del | 73 | 3 | 5.9E-07 | 9/28 | 10/34 | 2 |
| 4q16.3 | chr4:1705715-2073645 | Del | 75 | 0 | 2.6E-10 | 6/11 | 1/4 | 2 |
| 6p25.3 | chr6:259528-339802 | Del | 91 | 0 | 2.0E-12 | 0/1 | 0/0 | 0 |
| 6p25.3 | chr6:259528-339802 | Dup | 92 | 2 | 2.1E-12 | 0/1 | 0/0 | 0 |
| 7q11.23 | chr7:72742064-74142064 | Del | 146 | 3 | 1.5E-15 | 8/26 | 8/25 | 6 |
| 7q11.23 | chr7:72742064-74142064 | Dup | 108 | 5 | 1.5E-11 | 8/26 | 8/25 | 6 |
| 8p23.1 | chr8:10657597-10695288 | Del | 45 | 0 | 2.7E-06 | 0/3 | 0/0 | 0 |
| 8p23.1 | chr8:7268819-7366446* | Del | 54 | 1 | 2.9E-06 | 0/3 | 0/0 | 0 |
| 8p23.1 | chr8:7721060-7752586 | Dup | 50 | 0 | 4.5E-08 | 0/1 | 0/0 | 0 |
| 8p23.1 | chr8:8650757-11629240 | Dup | 72 | 2 | 4.1E-09 | 9/21 | 21/41 | 0 |
| 9q34.3 | chr9:140441351-140936261 | Del | 58 | 0 | 5.8E-08 | 2/8 | 2/4 | 1 |
| 10q23.1 | chr10:81960020-88800020 | Del | 118 | 15 | 3.9E-05 | 14/26 | 10/32 | 3 |
| 11p14.1 | chr11:29635361-31653568 | Del | 69 | 5 | 4.5E-05 | 8/8 | 9/11 | 1 |
| 13q34 | chr13:111699708-115085141 | Del | 66 | 0 | 5.2E-09 | 15/42 | 14/42 | 3 |
| 15q11.2 | chr15:22798636-23088559 | Del | 245 | 32 | 4.2E-09 | 2/4 | 0/5 | 0 |
| 15q11-13.1 | chr15:23094431-29410239 | Dup | 280 | 23 | 2.8E-21 | 14/17 | 28/47 | 5 |
| 15q11-13.1 | chr15:24674739-28514614 | Del | 147 | 19 | 6.6E-06 | 9/10 | 23/35 | 3 |
| 15q13.3 | chr15:30938215-32510863 | Del | 261 | 24 | 6.3E-14 | 5/11 | 4/13 | 2 |
| 15q13.3 | chr15:32452709-32509897 | Dup | 90 | 7 | 7.9E-08 | 1/1 | 0/0 | 1 |
| 16p13.11 | chr16:15248707-16292499 | Del | 171 | 26 | 1.8E-05 | 4/10 | 5/11 | 0 |
| 16p13.11 | chr16:15502499-16292499 | Dup | 288 | 64 | 8.6E-07 | 4/10 | 3/9 | 0 |
| 16p12.2 | chr16:21942499-22462224 | Del | 111 | 5 | 2.58E-12 | 4/8 | 3/6 | 0 |
| 16p11.2 | chr16:28822499-29052499 | Del | 81 | 2 | 9.4E-09 | 1/9 | 0/8 | 0 |
| 16p11.2 | chr16:29517499-30202499 | Dup | 275 | 28 | 5.8E-18 | 16/32 | 3/16 | 6 |
| 16p11.2 | chr16:29517499-30367499 | Del | 373 | 16 | 8.1E-31 | 16/34 | 3/21 | 6 |
| 17p13.3 | chr17:1038189-1673212 | Dup | 100 | 2 | 4.5E-13 | 6/22 | 2/17 | 4 |
| 17p11.2 | chr17:16446387-18380166 | Dup | 87 | 0 | 8.5E-14 | 17/36 | 12/26 | 5 |
| 17p11.2 | chr17:16789275-18299275 | Del | 61 | 1 | 2.7E-07 | 13/31 | 6/23 | 5 |
| 17p13.3 | chr17:2302359-3502250 | Dup | 85 | 5 | 1.9E-08 | 10/13 | 7/12 | 3 |
| 17q12 | chr17:34815887-34856055 | Dup | 86 | 6 | 6.2E-08 | 0/3 | 0/0 | 0 |
| 17q12 | chr17:34815887-36225887 | Del | 117 | 5 | 2.4E-10 | 4/17 | 5/12 | 1 |
| 17q21.31 | chr17:43704217-44164182 | Del | 86 | 0 | 8.9E-12 | 3/4 | 5/7 | 3 |
| 17p13.3 | chr17:911295-2593250 | Del | 65 | 3 | 5.1E-06 | 15/39 | 10/39 | 8 |
| 22q11.21 | chr22:18545000-21745000 | Del | 456 | 9 | 2.3E-47 | 18/54 | 10/37 | 5 |
| 22q11.21 | chr22:19020000-20308799 | Dup | 193 | 25 | 1.6E-10 | 6/32 | 5/17 | 3 |
| 22q11.22 | chr22:21910000-23650000 | Del | 89 | 9 | 6.3E-05 | 10/64 | 6/22 | 3 |
| 22q11.22 | chr22:23046186-23134119 | Dup | 46 | 0 | 1.2E-07 | 0/7 | 0/5 | 0 |
| 22q13.2-33 | chr22:43000056-51224208 | Del | 259 | 40 | 1.6E-07 | 45/113 | 31/109 | 8 |
| Xq28 | chrX:152561384-154917042 | Dup | 127 | 14 | 2.5E-08 | 17/70 | 4/19 | 6 |
| Xp22.33 | chrX:619146-2700156 | Dup | 146 | 22 | 6.0E-07 | 6/15 | 7/11 | 0 |
| Xp22.31 | chrX:6490000-7885155 | Del | 80 | 7 | 3.8E-05 | 1/4 | 1/4 | 0 |

\* chr8:7268819-7366446 is only significant when cohort (28) is kept.

Using a similar strategy to above we also tested whether the 118 significant regions were specific to individual cohorts, we removed case and controls from the five largest cohorts within AutDB and recalculated the *P* values for each region with the remainder of case and control samples. We repeated this analysis for each of the 5 cohorts separately. From this analysis we identified 12 regions that lost significance upon removal of one or more cohorts, which corresponds to 5 of the 47 independent regions mentioned above (**Supplementary table S7**). We suspect this is due to a reduction in power rather than a bias from a specific cohort as 9 of these lost significance by removal of the largest study (28) which itself is a collection of multiple geographically distinct cohorts.

### Potential ASD causal genes in regions identified using SNATCNV

For each of the 47 independent regions we next carried out literature searches to determine whether they contained known ASD genes or overlapped CNVs previously associated with ASD or intellectual disability (summarised in **Supplementary tables S8 and S9**). As expected, many genes from these independent regions (e.g. *NRXN1, GABRB3, MBD5, SHANK3, and MECP2*) show association with ASD (29–32). In total, 17 regions contained at least one gene that is known to be mutated in non-CNV mediated ASD (34 known ASD genes in 17 regions). Comparison of the 47 independent regions with CNV regions in DECIPHER, SFARI and MSSNG confirmed 34 overlapped previously identified syndromic/pathogenic regions with ASD-related neurodevelopmental phenotypes (e.g. autism, intellectual disability, delayed speech and language development, global developmental delay). Further systematic literature searches confirmed three of the remaining regions had strong evidence of ASD association (33–37) and a further seven were possibly associated with intellectual disability, but not specifically ASD (**Supplementary table 9**). Together this analysis highlights that the regions identified by SNATCNV are significantly associated with autism and intellectual disability.

For the 30 regions without a known causal gene we next asked whether any of the brain-enriched genes were strong candidates to cause ASD phenotypes. 22 of the 30 regions contained at least one brain-enriched coding gene (13 of them contained the human homologs of genes associated with mouse nervous system phenotypes). As an example, a deleted super region on chromosome 13 (chr13:111699708-115085141) contains no known ASD gene. However, it contains 15 brain-enriched coding and 14 brain-enriched lncRNA genes. Of note, mutations in mouse orthologs of three of the brain-enriched coding genes (*SOX1*, *ATP11A*, *GRK1)* cause nervous system phenotypes. Similarly, independent region chr17:1038189-1673212 contains no known ASD genes but contains six brain-enriched coding genes and two brain-enriched lncRNAs. Of these, mutations in mouse orthologs of four of the brain-enriched coding genes (*ABR*, *YWHAE*, *CRK, PITPNA)* cause nervous system phenotypes. Interestingly, one of the remaining 8 regions (chr2:242930600-243007359) lacks coding genes but does contain a novel brain-enriched lncRNA (ENSG00000237940), suggesting it may be important in ASD (**Supplementary figure S10**.

### Chromosomal rearrangements in males and female ASD patients

Given the 4-fold higher incidence of autism in males compared to females (38), we next examined the frequency of the 47 regions in a focused analysis on 8,083 cases (6,271 males and 1,812 females) and 2,741 control individuals (1,289 males and 1,452 females) where gender information was available. 22% of male and 37% of female cases had a CNV overlapping at least one of the 47 regions. When we compared the ratios of cases to controls for males and females, we identified 2 regions that were at least 2-fold more likely in male cases and 13 regions that were at least 2-fold more likely in female cases (**Table 2** and **Supplementary table S10** for more details). While further analyses will be needed to confirm our results, we note that one of the male-biased regions (chr16:15248707-16292499) and one of the female-biased regions (chr17:16789275-18299275) that we identify have been previously reported as gender biased (39, 40).

**Table 2.**
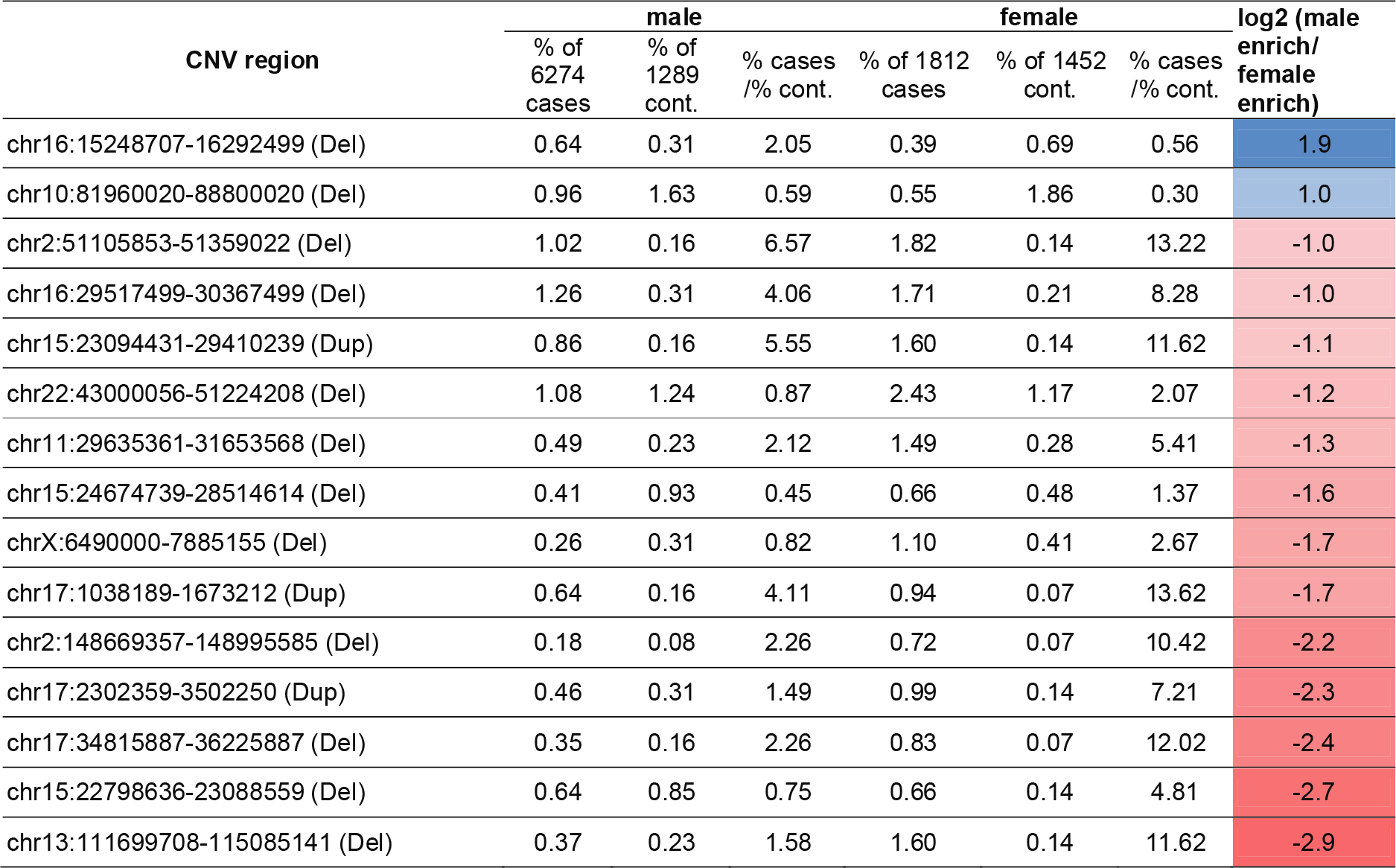
Sex bias of recurrent ASD associated CNV regions. Blue = male enriched, red = female enriched. Only regions that explain at least 0.6% of male or female cases and with at least 2 fold sex bias are shown. Extended statistics for all 47 regions are provided in **Supplementary table S10**.

| CNV region | male |  |  | female |  |  | log2 (male enrich/<br>female enrich) |
| --- | --- | --- | --- | --- | --- | --- | --- |
|  | % of 6274 cases | % of 1289 cont. | % cases /% cont. | % of 1812 cases | % of 1452 cont. | % cases /% cont. |  |
| chr16:15248707-16292499 (Del) | 0.64 | 0.31 | 2.05 | 0.39 | 0.69 | 0.56 | 1.9 |
| chr10:81960020-88800020 (Del) | 0.96 | 1.63 | 0.59 | 0.55 | 1.86 | 0.30 | 1.0 |
| chr2:51105853-51359022 (Del) | 1.02 | 0.16 | 6.57 | 1.82 | 0.14 | 13.22 | -1.0 |
| chr16:29517499-30367499 (Del) | 1.26 | 0.31 | 4.06 | 1.71 | 0.21 | 8.28 | -1.0 |
| chr15:23094431-29410239 (Dup) | 0.86 | 0.16 | 5.55 | 1.60 | 0.14 | 11.62 | -1.1 |
| chr22:43000056-51224208 (Del) | 1.08 | 1.24 | 0.87 | 2.43 | 1.17 | 2.07 | -1.2 |
| chr11:29635361-31653568 (Del) | 0.49 | 0.23 | 2.12 | 1.49 | 0.28 | 5.41 | -1.3 |
| chr15:24674739-28514614 (Del) | 0.41 | 0.93 | 0.45 | 0.66 | 0.48 | 1.37 | -1.6 |
| chrX:6490000-7885155 (Del) | 0.26 | 0.31 | 0.82 | 1.10 | 0.41 | 2.67 | -1.7 |
| chr17:1038189-1673212 (Dup) | 0.64 | 0.16 | 4.11 | 0.94 | 0.07 | 13.62 | -1.7 |
| chr2:148669357-148995585 (Del) | 0.18 | 0.08 | 2.26 | 0.72 | 0.07 | 10.42 | -2.2 |
| chr17:2302359-3502250 (Dup) | 0.46 | 0.31 | 1.49 | 0.99 | 0.14 | 7.21 | -2.3 |
| chr17:34815887-36225887 (Del) | 0.35 | 0.16 | 2.26 | 0.83 | 0.07 | 12.02 | -2.4 |
| chr15:22798636-23088559 (Del) | 0.64 | 0.85 | 0.75 | 0.66 | 0.14 | 4.81 | -2.7 |
| chr13:111699708-115085141 (Del) | 0.37 | 0.23 | 1.58 | 1.60 | 0.14 | 11.62 | -2.9 |

### Specific phenotypes associated with distinct regions of copy number variations

We next used the DECIPHER (17) database, containing detailed phenotypic information for over 12,795 patients with CNVs, to identify patient phenotypes specifically associated with each of the refined 47 SNATCNV-identified ASD CNV regions. Given that DECIPHER contains 1,360 patients annotated as autistic, 5,559 patients annotated with intellectual disability (637 both annotated as ASD and ID) and 6,513 patients with other phenotypes, we quantified the number of patients which overlapped each one of the 47 independent CNV regions and compared it to the total number of patients in DECIPHER annotated with distinct Human Phenotype Ontology (HPO) terms. At 95% confidence interval 34 of the 47 super regions had at least one phenotypic term that was 2-fold over-represented in DECIPHER patients with CNVs overlapping the region (**Figure 3**, **Supplementary table S11**).

**Figure 3.**
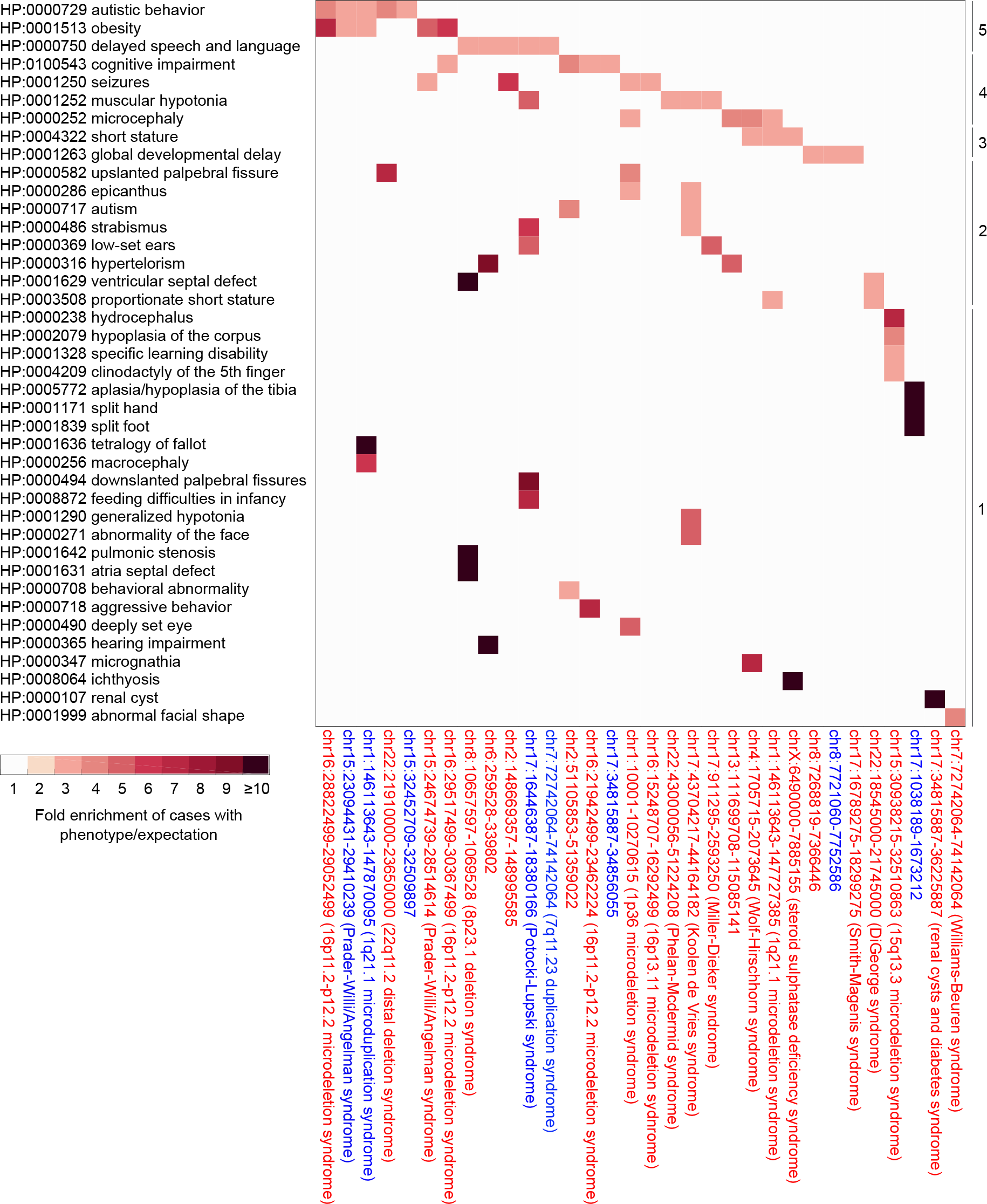
Association of specific patient phenotypes recorded in the DECIPHER database for 34 of the 47 super regions (indicated on the X-axis). Overrepresented human phenotype ontology terms are indicated by red shaded boxes. Here, we only report region-phenotype term pairs that are at least 2 fold overrepresented, present in at least 5 patients, seen in at least 5% of patients in the region and at 95% C.I. (also see **Supplementary table S11**). Regions associated with known syndromes are indicated by descriptions in brackets.

The most commonly overrepresented terms were HP:0000729 autistic behaviour, HP:0001513 obesity and HP:0000750 delayed speech and language development, each found overrepresented in 5 of the SNATCNV-identified ASD CNV regions. The next most common terms were HP:0100543 cognitive impairment, HP:0001250 seizures, HP:0001252 muscular hypotonia, and HP:0000252 microcephaly. Of note, three regions had enrichment for both autistic behaviour and obesity. chr16:28822499-29052499 (16p11.2) has previously been associated with ASD and obesity (41) and chr15:23094431-29410239 (15q11-12), corresponds to the Prader-Willi syndromic region (8). The remaining region, deletion of chr1:146113643-147870095 (1q21.1-21.2) is not only associated with HP:0000729 autistic behaviour and HP:0001513 obesity but also HP:0001636 tetralogy of fallot and HP:0000256 macrocephaly. Interestingly duplication of the overlapping region chr1:146113643-147727385 is associated with the reciprocal phenotype HP:0000252 microcephaly (42), which could suggest dosage sensitivity of constituent genes within the loci. Three brain-enriched coding genes *NBPF12*, *FMO5*, *BCL9* and two brain-enriched lncRNA genes lie within a region of overlap between the deleted and duplicated regions, implicating these as candidate genes in microcephaly and macrocephaly.

Of note, half of the CNV regions of interest (23/47) had only one overrepresented term. For example, HP:0004209 clinodactyly of the 5th finger was overrepresented in patients with deletion of chr15:30938215-32510863 (2.5-fold) while HP:0001631 atria septal defect was overrepresented for patients with deletions of chr8:10657597-10695288 (24 fold). As another example, HP:0008064 ichthyosis was overrepresented in patients with deletion of chrX:6490000-7885155 (64-fold). Steroid sulfatase gene (STS) is the only brain-enriched gene in this region and it has been reported that X-linked ichthyosis caused by deletions or point mutations of the STS gene are associated with increased risk of ASD (43).

In addition to the identification of CNV regions associated with unique phenotypes we also found evidence of segmentation of single syndromic regions into multiple, distinct sub-loci defined by unique phenotypic traits. For example, three of our deleted regions overlap the known syndromic deletion region 16p11.2-p12.2. Note in the literature, sub-loci in this region have been previously described based on the presence of 5 distinct breakpoints (BP1-BP5 (44)). Two of the regions we identify, chr16:28822499-29052499 and chr16:29517499-30367499 correspond to BP2-BP3 and BP4-BP5 respectively, however chr16:21942499-22462224 is outside of BP1-BP5 (see **Supplementary figure S11**). Examining the phenotypes we note, chr16:21942499-22462224 is associated with HP:0100543 cognitive impairment and HP:0000718 aggressive behaviour, chr16:28822499-29052499 is associated with HP:0000729 autistic behaviour and HP:0001513 obesity; and chr16:29517499-30367499 is associated with HP:0001513 obesity and HP:0100543 cognitive impairment. Based on our analysis, SNATCNV is able to identify these as separate regions and the DECIPHER analysis demonstrates they are associated with different phenotypes.

Taken together, our implementation of SNATCNV has led to the curation of ASD CNVs which are enriched in genes expressed in the nervous system, including lncRNAs; and that the diversity of phenotypic traits in ASD caused by CNVs can likely be defined by unique genomic signatures.

## Discussion

We developed a new computational method, SNATCNV, to systematically analyse recurrent copy number variations and applied it to ASD. It equally can be applied to any genetic disease and disorder with sufficient cases and controls. SNATCNV more precisely identifies CNV regions, and more accurately discriminates cases from controls, thereby reducing the genomic search space for ASD and its causal genes. We applied SNATCNV to investigate 19,663 autistic and 6,479 healthy individuals and found 118 CNV regions that contain an overrepresentation of brain-enriched coding and lncRNA genes. The 118 regions were then further narrowed to 47 CNV super regions by merging co-dependent regions and removing co-deleted and co-duplicated (dependent) regions. Of the 47 regions, 10 overlapped SFARI syndromic regions, 24 overlapped DECIPHER neurodevelopmental related syndromic regions (see methods), and 36 overlapped with regions previously associated with developmental delay (20, 21). Interestingly, 22 of the 47 regions overlapped with CNV regions recently identified in whole genome sequencing carried out by MSSNG (5). Our systematic literature search also revealed that 17 of the 47 regions contained at least one known ASD causal gene with recorded single gene mutations from MSSNG, SFARI and other studies. For some of these regions, we report additional ASD candidate genes, which based on their brain-enriched expression are more likely candidates. For example, in the duplicated region “chr22:19020000-20308799”, there are two genes where an association between rare variants and ASD (i.e. *CLTCL1* (45), *GNB1L* (46)) have been made. However, neither of these are brain-enriched and both are classified by SFARI gene as category 4 ‘Minimal Evidence’ ASD candidate genes. We identify six brain-enriched coding genes in this region. Mutation of mouse orthologs of three of these (*SEPT5*, *ZDHHC8*, and *RTN4R*) cause nervous system phenotypes suggesting these genes are better potential candidates for this region.

Supporting this hypothesis that brain-enriched genes are more probable ASD casual genes than others we note that 63% of high-confidence ASD genes from MSSNG and SFARI consortiums are brain-enriched (**Fig. 2b**). Thus for the 29 regions without a known ASD causal gene we provide lists of brain-enriched ASD candidate genes for further investigation. As an example, in deleted region chr17:43704217-44164182, we identify three brain-enriched coding genes *MAPT*, *CRHR1* and *KANSL1*. All three cause nervous system phenotypes in mutant mice, and all three have been linked to neurological phenotypes in human (deletion of *MAPT* is associated with developmental delay and learning disability (47), *CRHR1* controls stress-induced behavioural adaptations and is associated with mood disorders (48, 49) and *KANSL1* deletion in 17q21.31 deletion syndrome is associated with intellectual disability, hypotonia and friendly behaviour (50)). It is interesting to hypothesize that loss of each individually may be insufficient to cause ASD but the loss of all three contributes to an ASD phenotype. Totally, we report 299 novel ASD candidate coding genes.

In addition to brain-enriched coding genes we also report for the first time to our knowledge, 266 novel recurrently duplicated or deleted ASD candidate lncRNAs, which are brain-enriched. Suggesting an importance in neurodevelopment, 94 are dynamically expressed in an induced pluripotent stem cell model of neuron differentiation (**Supplementary table S8**). We note that although lncRNAs that are differentially expressed in post mortem brain of ASD cases have been reported there is no evidence of them playing a role in ASD causation (51). In contrast the lncRNAs we identify here are in recurrent CNV regions, are brain-enriched and in the case of one lncRNA (ENSG00000237940) is the only gene in the region. This suggests that this lncRNA is the most likely ASD candidate gene for this region.

In establishing phenotypic associations for the 47 super CNV regions, our analysis using DECIPHER classification terms identified overrepresented traits including autistic behaviour, delayed speech and language, cognitive impairment and global developmental delay. For 34 of the 47 identified regions, at least one phenotype term was over-represented (fold enrichment above 2). Of note the majority of these 34 regions had unique combinations of overrepresented phenotypes (40 combinations in total), thus we conclude that ASD CNV regions are associated with distinct clinical phenotypes. The observation of these combinations of additional phenotypes may assist in diagnosis of patients (e.g. obesity, microcephaly, macrocephaly, split hand). Importantly, for patients with a subset of these CNVs, genomic diagnosis will inform the clinician that these patients should be examined and potentially treated for associated co-morbidities such as renal cysts, atria septal defects and tetralogy of fallot.

In conclusion by application of a novel tool SNATCNV, integration of brain-enriched expression profiles from the FANTOM5 consortium and phenotypic information from DECIPHER we have annotated 47 recurrent CNV regions associated with ASD. Looking to the future, screening for these CNVs and single gene variants identified by the MSSNG and SFARI consortia using whole genome sequencing has the potential to provide a rapid and accurate genetic diagnosis for children with ASD. By knowing the genes and phenotypes associated with the region a personalised treatment plan can be developed that considers the specific phenotypes associated with their genomic lesion.

## Materials and methods

### Tool availability

The SNATCNV source code for both R and matlab versions, a sample dataset, and instructions on how to run SNATCNV are provided at https://github.com/hamidrokny/SNATCNV.

### ASD and control data used in this study

For the region discovery, CNV data from 19,663 autistic patients (47,189 CNVs) and 6,479 normal individuals (24,888 CNVs) compiled from multiple large ASD CNV datasets (with different microarray versions and different ‘individual-level’ CNV callers) by the Simons Foundation Autism Research Initiative (SFARI) was downloaded (http://autism.mindspec.org/autdb; download date: July 2016).

For the validation cohorts, CNV data from i) 1,441 autistic patients were downloaded from the same source (download date July 2018), and ii) 549 autistic individuals were downloaded from DECIPHER (download date 1 Feb 2017; we excluded those individuals that were also annotated with intellectual disability, developmental delay, heart and muscular diseases). As CNVs from different studies were reported using different genome build versions (hg17, hg18, and hg19), we first converted all CNV coordinates to UCSC hg19 using UCSC Lift Genome Annotations tools (52) and confirmed the locations with NCBI remap tools (*www.ncbi.nlm.nih.gov/genome/tools/remap*). We excluded all CNVs smaller than 1kb or where there was no information provided about their genomic coordinates. In the sex specific analysis, we excluded those samples with no sex information.

### Genome-wide association analysis using SNATCNV to identify significant regions

To identify regions that are significantly more often deleted or duplicated in autism patients than in phenotypically normal individuals, we counted, for every position in the genome, how often the base was either deleted or duplicated in the case and control CNVs. Fisher’s exact test (**Supplementary figure S7**) was then used to determine whether the base was significantly more frequently observed in a duplicated or deleted region of cases or controls. To determine the significance of this association we then used permutation testing by randomising 500,000 permutations of case and control labels and for a range of observed CNV frequencies from 1-700 (min and max number of CNVs that overlapped any base in the original data). In each permutation, we randomly selected case/control labels and then estimated the probability of the observed associations using the Fisher’s exact test. By performing this permutation 500,000 times and for CNV frequency 1-700, we could estimate the probability that an association at a given level of CNV frequency emerges by chance. From this we defined significant regions as those with *P* values better than 1.2e−08 (approaching 100% confidence) (**Supplementary figure S8**). Note, due to the poor coverage of CNVs on chrY it was not included in the analysis.

### PLINK and ParseCNV comparisons

Identical AutDB sample sets were run on both SNATCNV and PLINK version 1.07 (http://zzz.bwh.harvard.edu/plink/download.shtml). We used the burden-style methods in PLINK “—cnv-indiv-perm” with 50,000 permutations and the default options in PLINK, which returns one-sided empirical *P* values, assuming that the regions are more common in case individuals than in control individuals.

As ParseCNV requires input files in PennCNV format we were unable to use the same dataset as above. Consequently the ParseCNV comparison was run using a CNV dataset provided on the ParseCNV website (link to data: https://sourceforge.net/projects/parsecnv/files/TestDataParseCNV_Plink_AGRE_CaseControl.tgz/download). CNV carrier frequencies between case and control individuals were compared using the one-tailed Fisher exact test and those regions with *P* value < 0.05 considered as significant regions. Both SNATCNV and PLINK were run on this PennCNV format dataset for direct comparison (**Supplementary figure S4**).

### Identification of genes with nervous system enriched expression

To determine whether a gene had nervous system enriched expression we first extracted its normalised expression profile from the FANTOM5 expression atlas (FANTOM5 (16) and FANTOMCAT (19)). We chose this resource as it contains expression profiles for both coding and long-non-coding RNA genes across 1,829 human samples, including 101 nervous system samples. For each gene we first ranked all 1,829 human FANTOM5 samples based on highest to lowest expression and then used Fisher’s exact test to determine a *P* value indicating whether the top 101 samples with the highest expression were more likely to be nervous system samples or not.

To determine a threshold on the *P* value we carried out 50,000 permutations randomising the sample labels to determine *P* value thresholds at the 99% confidence interval; any gene with a *P* value better than this permutation-specific *P* value threshold, we consider a significantly nervous system-enriched gene (**Supplementary table S3**). The same strategy was used to identify genes with enriched expression in blood, muscle, and sub-regions of the brain. A list of all brain-enriched genes is provided in **Supplementary table S12**.

### Identification of genes with mouse nervous system phenotypes

Mouse genes with nervous system phenotypes were extracted from the Mouse Genome Informatics (MGI) resource (http://www.informatics.jax.org; June 12^th^ 2018) (53). Specifically HTML was parsed from the following pages covering, (i) Nervous system phenotype (MP:0003631) http://www.informatics.jax.org/mp/annotations/MP:0003631, (ii) abnormal nervous system morphology (MP:0003632) http://www.informatics.jax.org/mp/annotations/MP:0003632 and (iii) abnormal nervous system physiology (MP:0003633) http://www.informatics.jax.org/mp/annotations/MP:0003633. Mouse to human gene ortholog mappings were obtained from http://www.informatics.jax.org/downloads/reports/HGNC_homologene.rpt.

### Gene ontology, disease ontology and human phenotype ontology analyses of nervous system enriched genes in the 118 regions in the brain candidate genes

We used the gene ontology analysis tool WebGestalt (54) to observe over-representation of gene ontology, disease ontology and human phenotype ontology terms annotations associated with our genes in the significant regions. Test gene set corresponded to all nervous system enriched coding genes found within the 118 regions. Background gene set corresponded to all nervous system enriched coding genes found in the entire genome. Default values for WebGestalt parameters were used.

### DECIPHER analysis

Phenotypic information of patients with copy number variation were taken from the publicly available DECIPHER database (17), which contains phenotypic information of ~13,000 patients (including 1360 patients annotated as autistic and 5,559 patients annotated with intellectual disability). To assess whether a phenotype term was over-represented for a specific region, we used 5,000 random permutations of the patient-phenotype associations in DECIPHER. In order to keep the data structure of the CNVs, we needed to permute all the phenotypes of a given patient *en bloc*. It means in each permutation, the patient details (e.g. CNV, sex) do not change but all the phenotypes will be changed. For each permutation, we had a new order of phenotype sets randomly matched to patient sets. By performing this permutation 5,000 times, we could estimate the fraction of patients with the same region-phenotype pair that emerges by chance. Because of the small number of samples in some regions, we only reported those region-phenotype term pairs that had a fraction more than the permutation-specific fraction and seen in at least 5% of patients with a minimum of 5 patients overlapping the region.

### Annotation of known and putative novel ASD associated genes

All genes in the SNATCNV identified regions were first compared to the 96 known causal genes from MSSNG [5] and SFARI [4, 15] (Supplementary table S13). If a region contained one or more of these genes we considered it ‘known’. For the remaining regions, we carried out a comprehensive literature search of all genes in the region. For the extended literature search, we specifically searched for “gene name + autism” or “gene name + ASD” (e.g. SAMD11 + autism or SAMD11 + ASD). We did this search for all coding genes in the 47 super regions. As a result, if a gene with a previous ASD casual association was identified it was labelled as ‘known’.

### Annotation of known and putative novel ASD associated CNVs

To investigate whether the regions we identified have been previously associated to ASD, we compared the regions identified by SNATCNV to the data from DECIPHER (17), SFARI [4, 15], MSSNG [5] and to the two largest CNV studies of Global developmental delay to date by Coe *et al.* [21] and Cooper *et al*. [22]. We also carried out a comprehensive literature search. As a result of our investigation, If the region completely overlap with a known ASD-associated CNV, it was labelled as “known CNV”; if the region contains a known ASD-associated CNV region (the known CNV was a subset of the region), it was labelled as “contains known CNV”; if the region was a subset of a known ASD-associated CNV, but smaller, it was labelled as “known CNV, but finer mapping”, otherwise, it was labelled as “novel”.

## Author contributions

HAR and ARRF designed the study and wrote and edited the manuscript with help from JI-TH. HAR carried out all the analyses including the statistical analyses, region identification, text mining, gene prioritization, and gene ontology (with help from ARRF). JI-TH, HAR and ARRF curated the genes and regions.

HAR generated all figures and tables. All authors have read and approved of the final version of the paper.

## Author information

The authors declare no competing financial interests.

## Data availability

The SNATCNV source code and a sample dataset can be accessed at *https://github.com/hamidrokny/SNATCNV*. For more details about each parameter, please visit the github page.

## Acknowledgements

This work was supported by a grant to JI-TH and ARRF from the Telethon-Perth Children’s Hospital Research Fund. ARRF and JI-TH are funded by grants from the National Health and Medical Research Council (Australia), and ARRF is supported by a Senior Cancer Research Fellowship from the Cancer Research Trust and funds raised by the Maca Ride to Conquer Cancer.

## Supplementary figure legends

**Figure S1.**
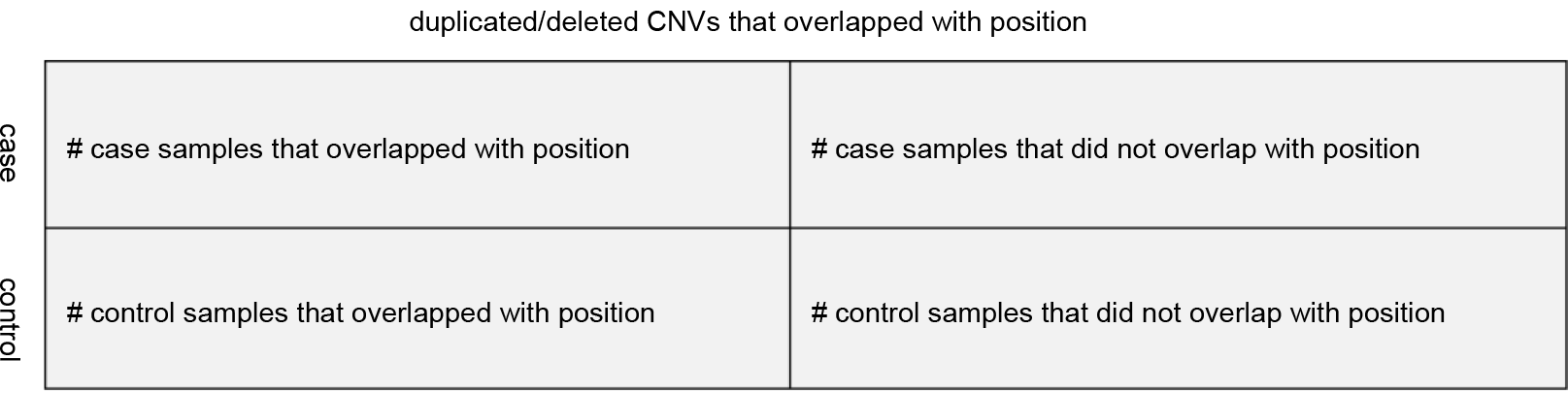
Contingency table for Fisher’s exact test to calculate *P* value for each genomic position. *P* value for deleted and duplicated regions are calculated separately.

**Figure S2.**
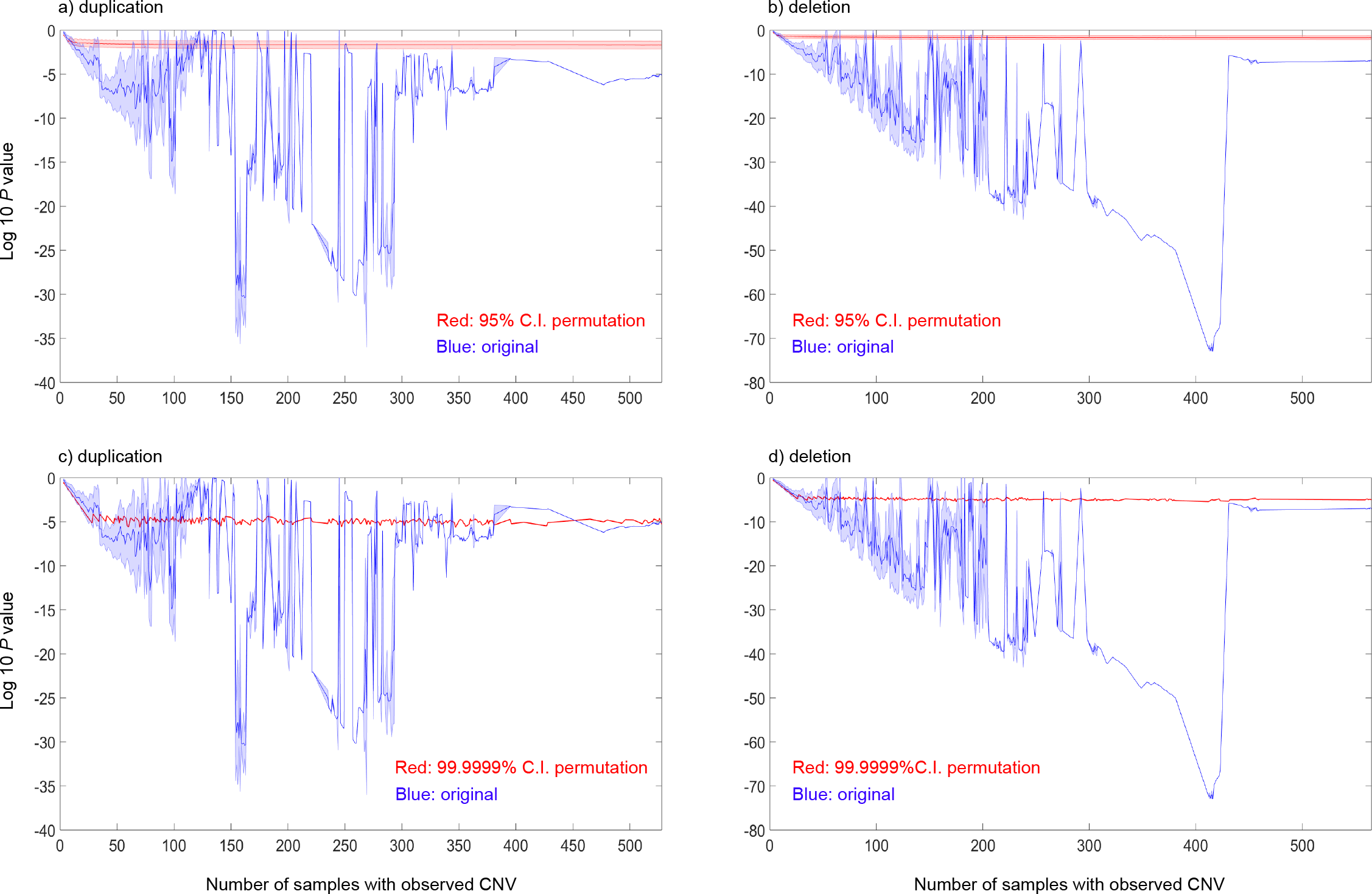
Permutation thresholds to identify significant regions at 95% and 99.9999% C.I. Blue colour indicates CNV signals from original data and red colour indicates CNV signals from permutated data. **a)** Duplication threshold at 95% C.I. **b)** Deletion threshold at 95% C.I. **c)** Duplication threshold at 99.9999% C.I. **d)** Deletion threshold at 99.9999% C.I. X-axis indicates number of individuals with observed CNV and y-axis indicates logarithm 10 *P* value. *P* value calculated through one-tailed Fisher exact test.

**Figure S3.**
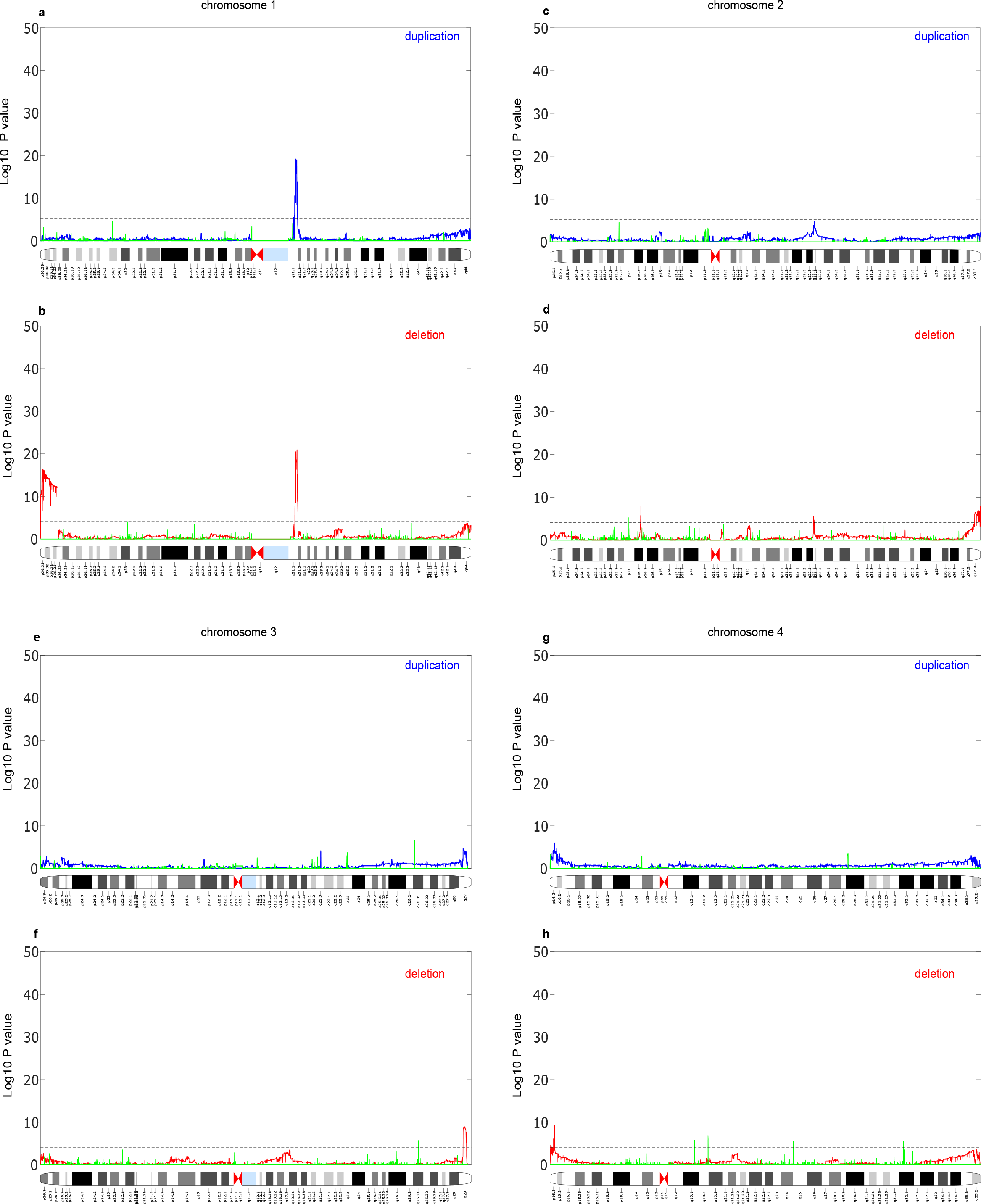
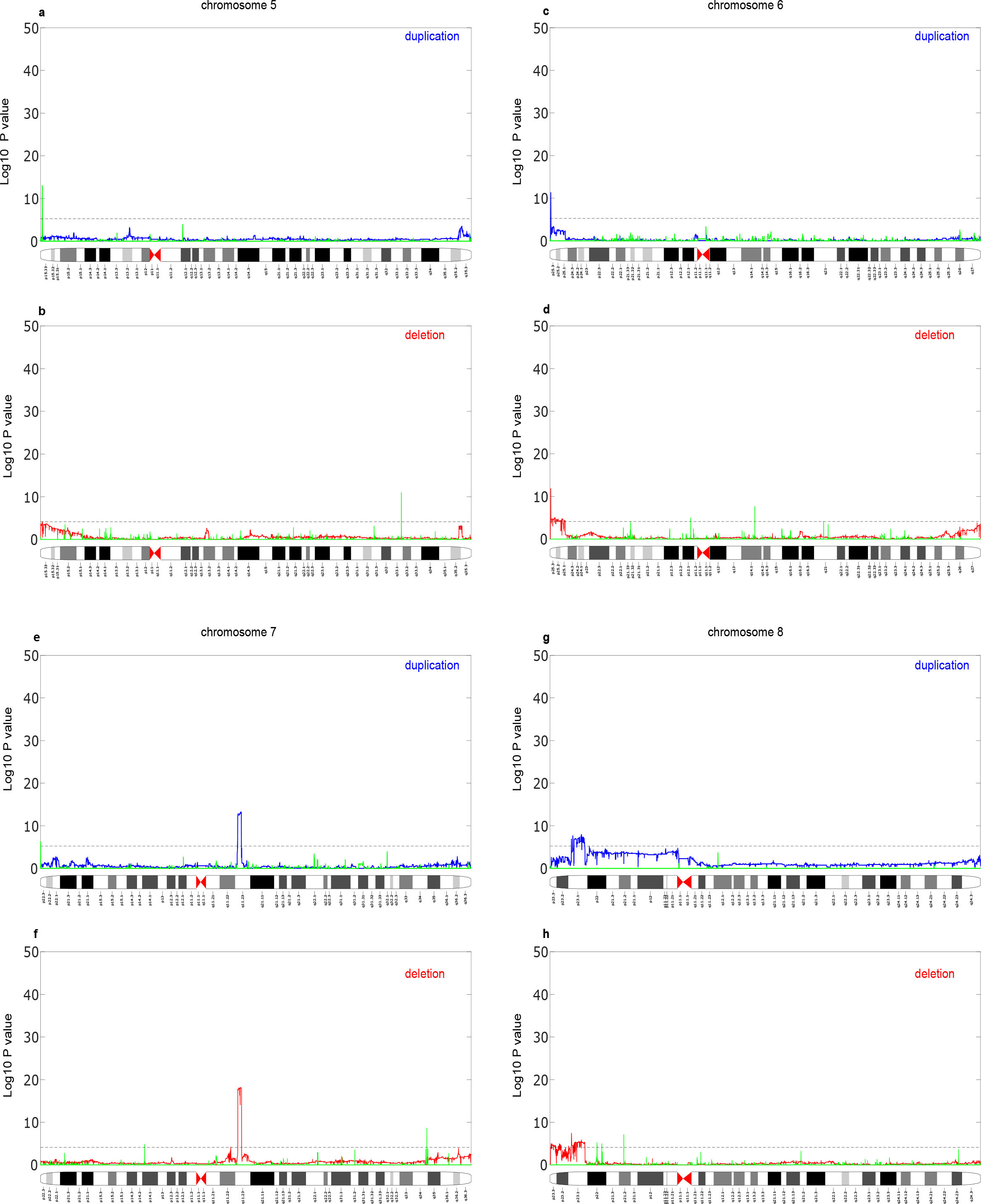
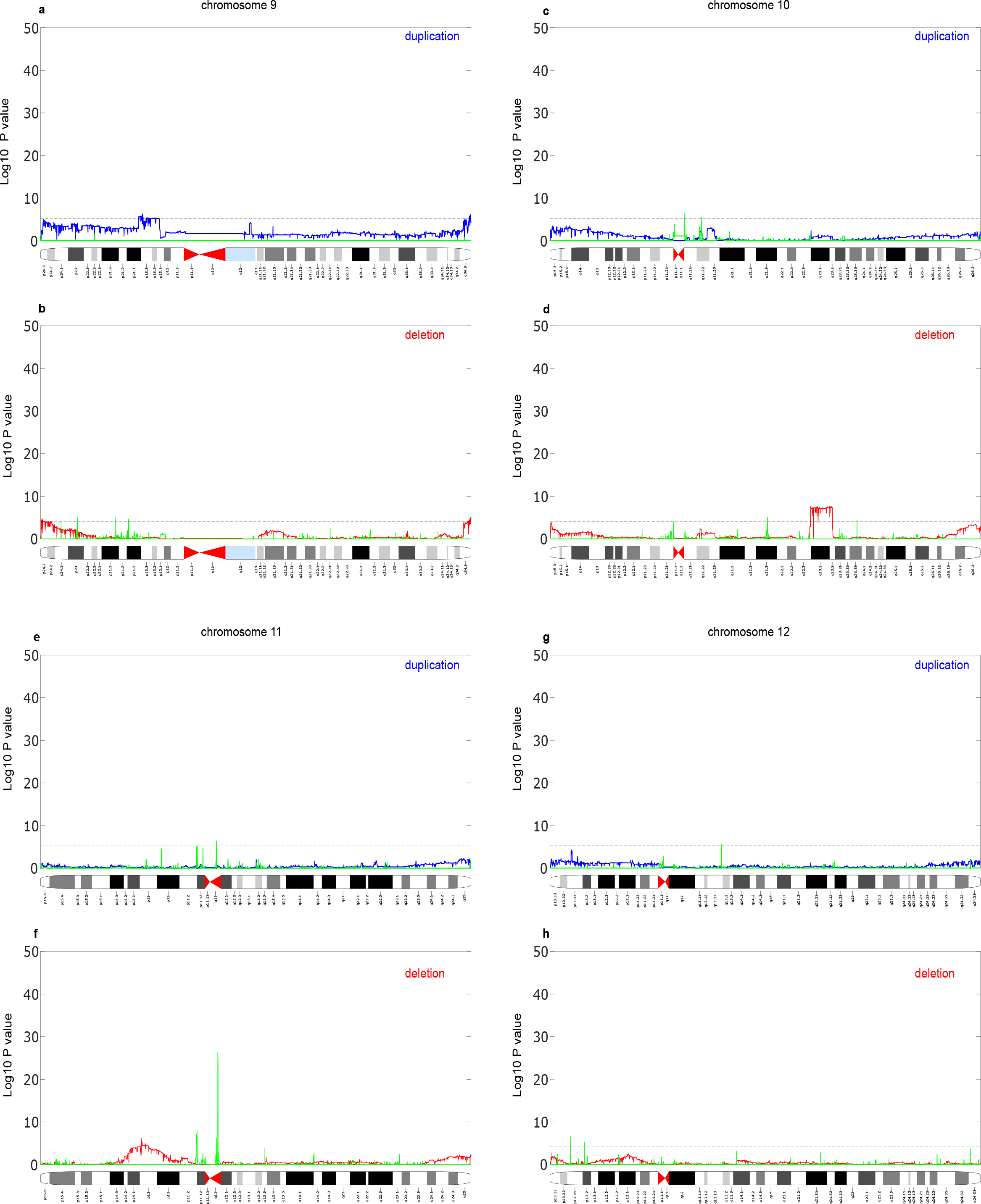
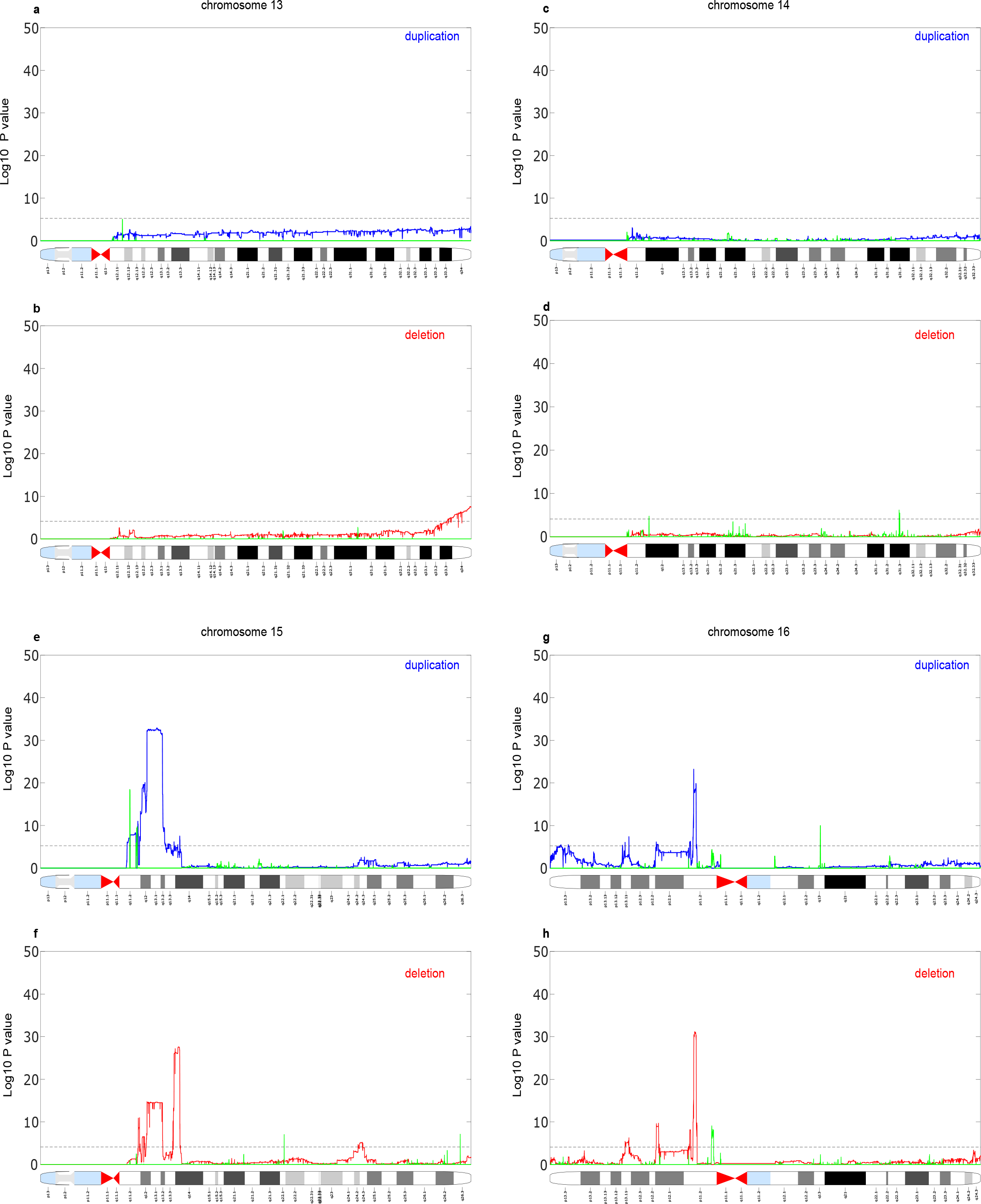
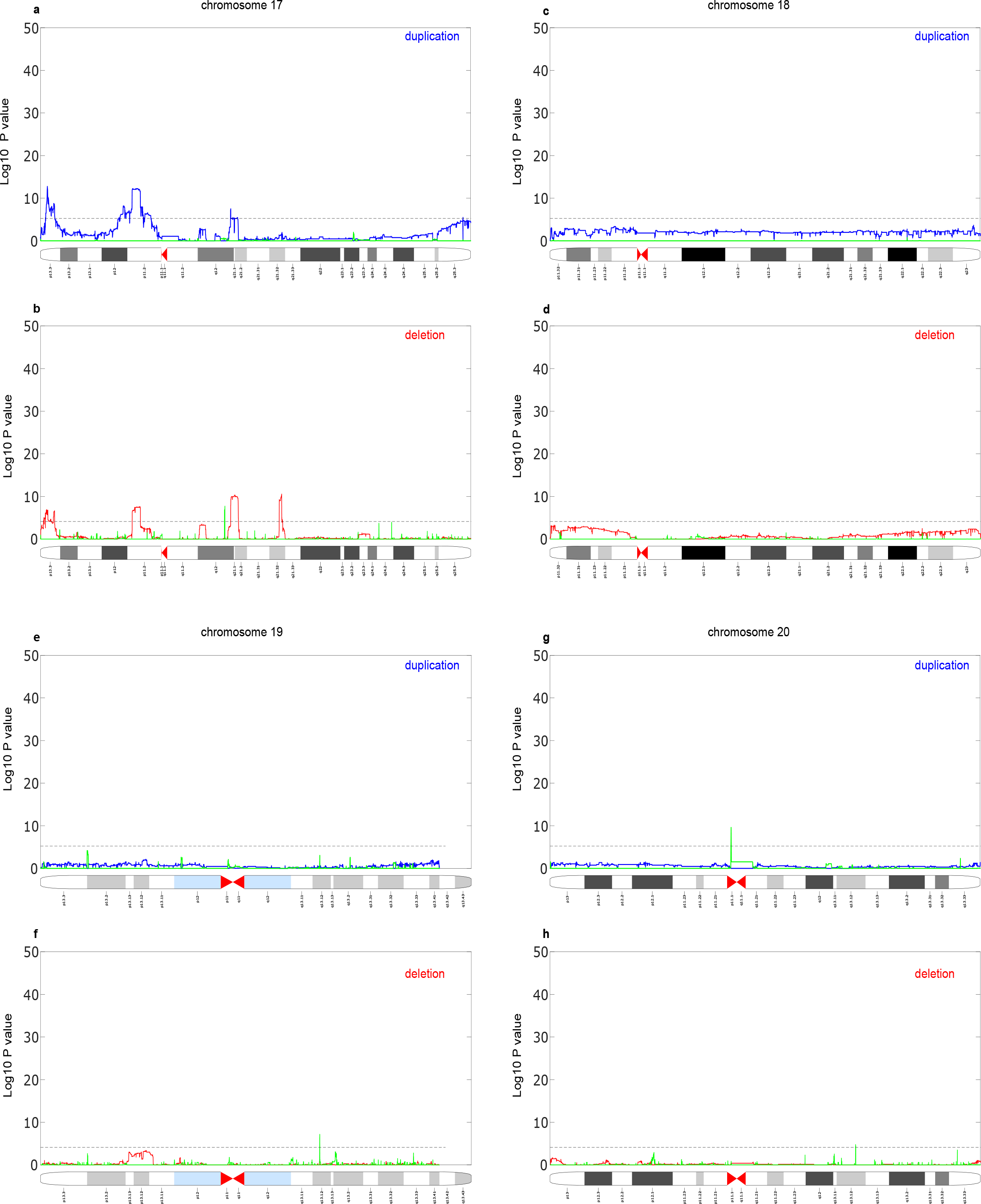
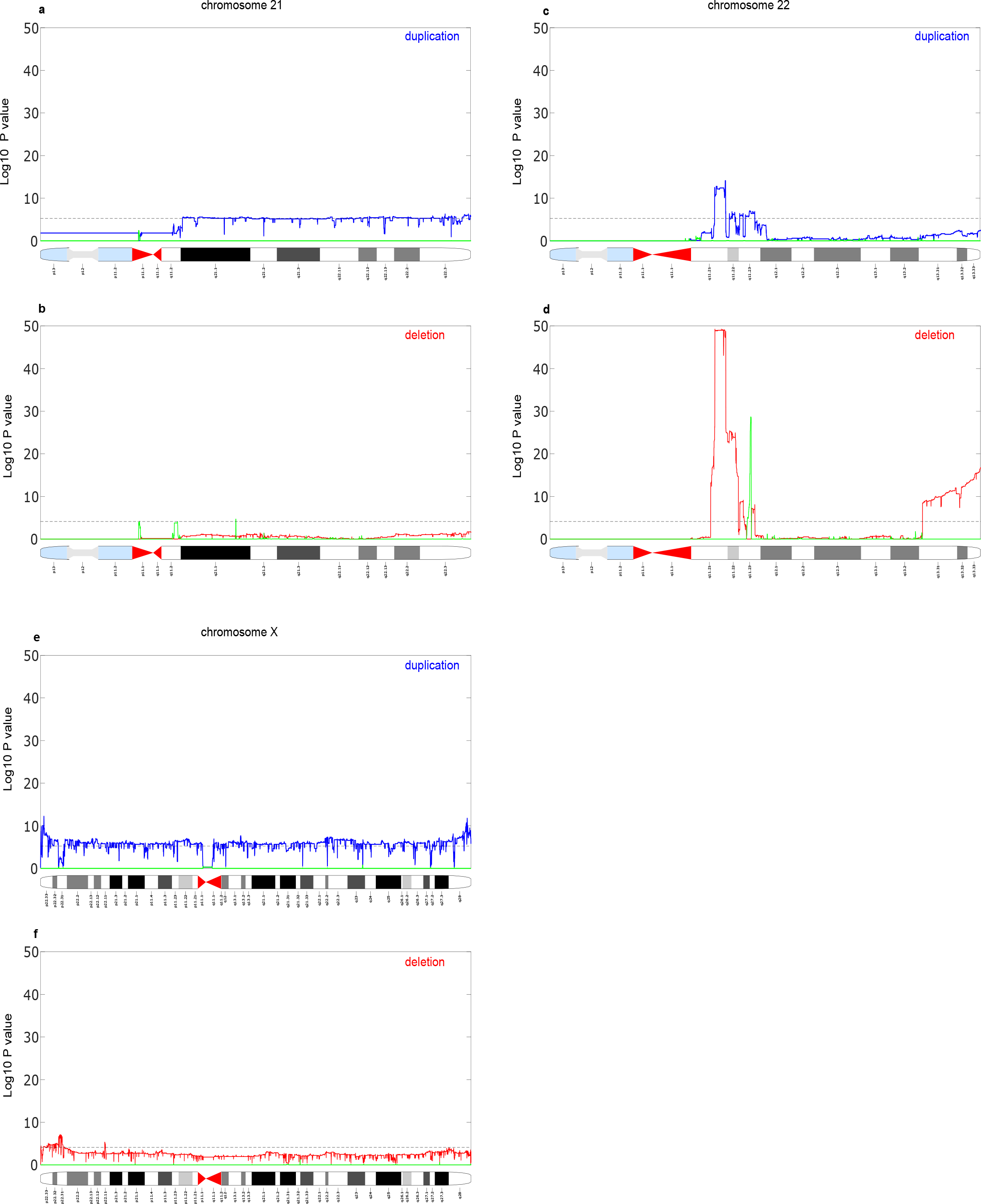
ASD copy number variation signals on human chromosomes. **a)** Autistic duplication vs control duplication copy number variation signals on chromosome 1; blue colour indicates autistic signals and green colour indicates control signals. **b)** Autistic deletion vs control deletion copy number variation signals on chromosome 1; red colour indicates autistic signals and green colour indicates control signals. Black dashed line indicates our threshold for significant regions (better *P* value than 99.9999 percentile of 500,000 permutations). X-axis indicates genomic loci and y-axis indicates logarithm 10 *P* value. *P* value calculated through one-tailed Fisher exact test. **c-h**) as in a, b but for the remaining chromosomes.

**Figure S4.**
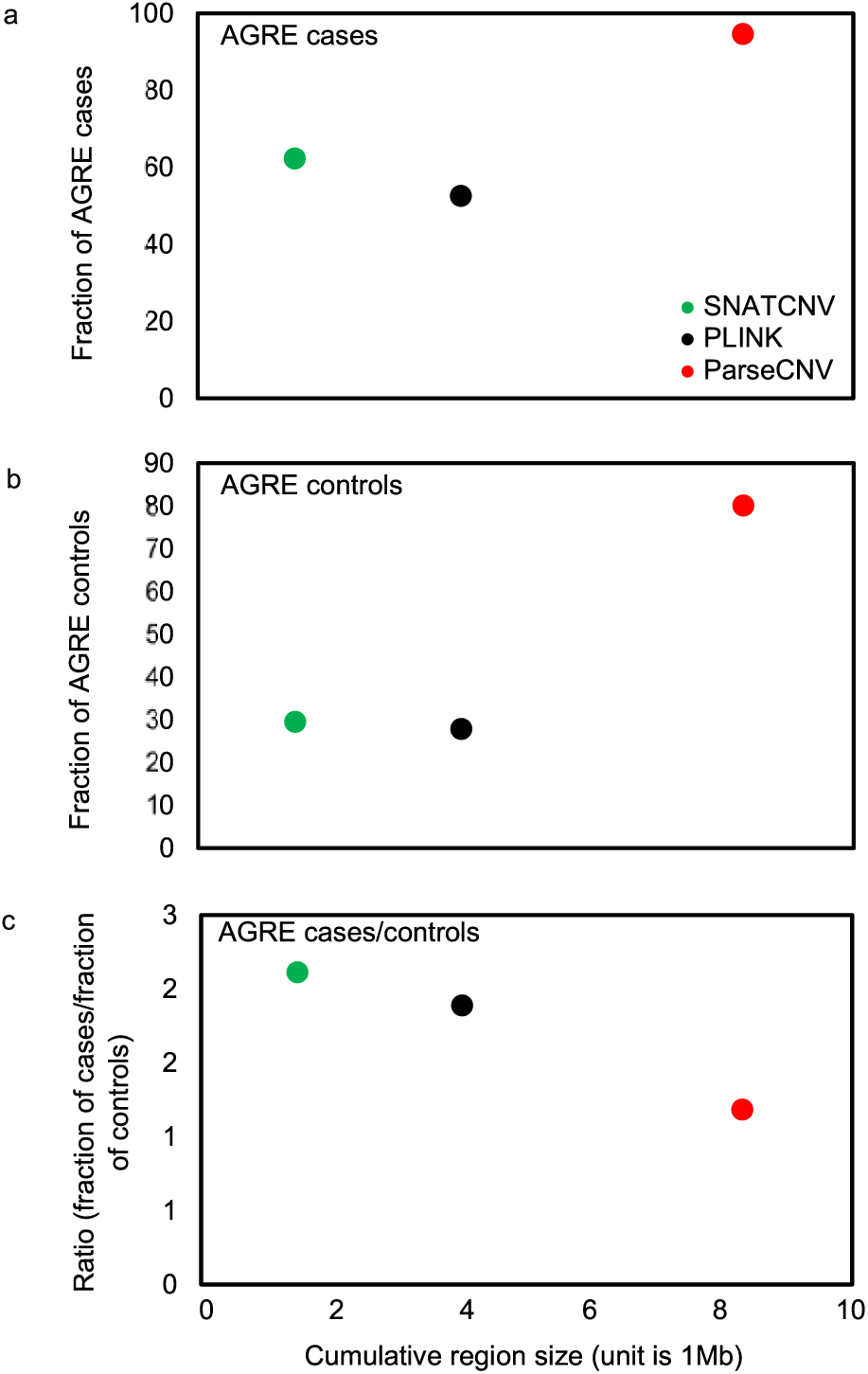
SNATCNV versus PLINK and ParseCNV on a PennCNV dataset. The plot shows the fraction of samples covered by SNATCNV’s regions versus PLINK and ParseCNV regions on a PennCNV dataset (http://parsecnv.sourceforge.net). Vertical axis is the fraction of ASD individuals and horizontal axis is region size. **a**) The plot shows the fraction of case samples covered by SNATCNV’s regions (green) versus those found by PLINK (black) and ParseCNV (red) **b)** The plot shows control samples covered by SNATCNV’s regions versus PLINK and ParseCNV regions. **c)** This plot shows the ratio of the fraction of ASD samples covered by the regions divided by fraction of control samples covered by the regions for SNATCNV, PLINK and ParseCNV.

**Figure S5.**
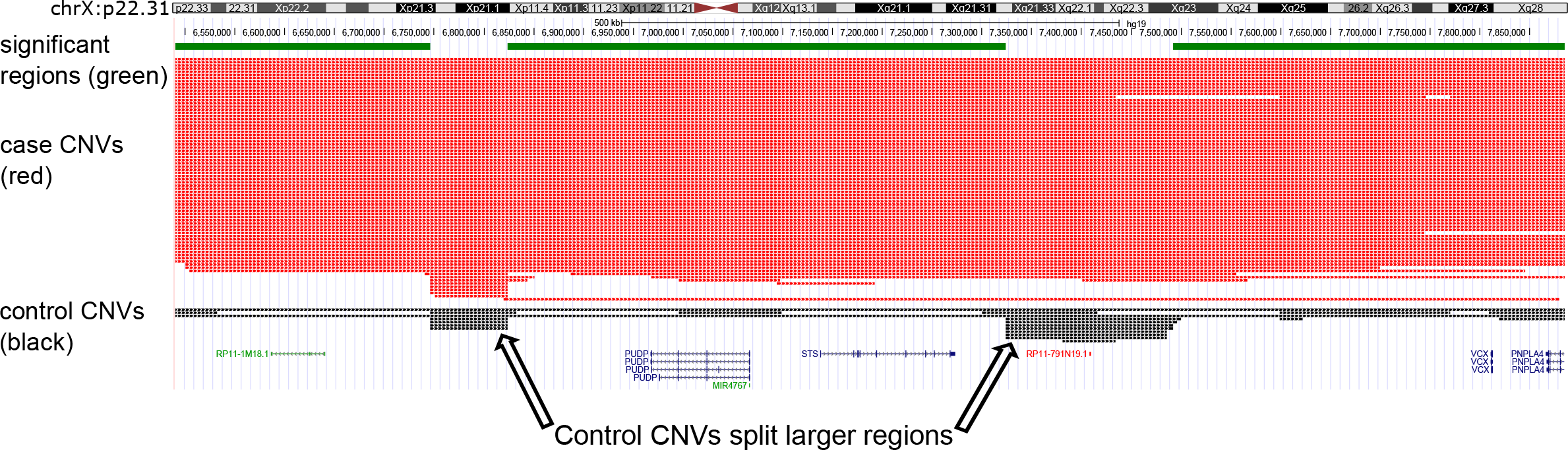
An overview of significant regions (chrX: 6,500,000-7,920,000) identified by SNATCNV split by control CNVs. Green colour indicates significant regions identified by SNATCNV; red colour indicates CNVs from autistic individuals; and black colour indicates CNV from healthy individuals. CNVs have different breakpoints.

**Figure S6.**
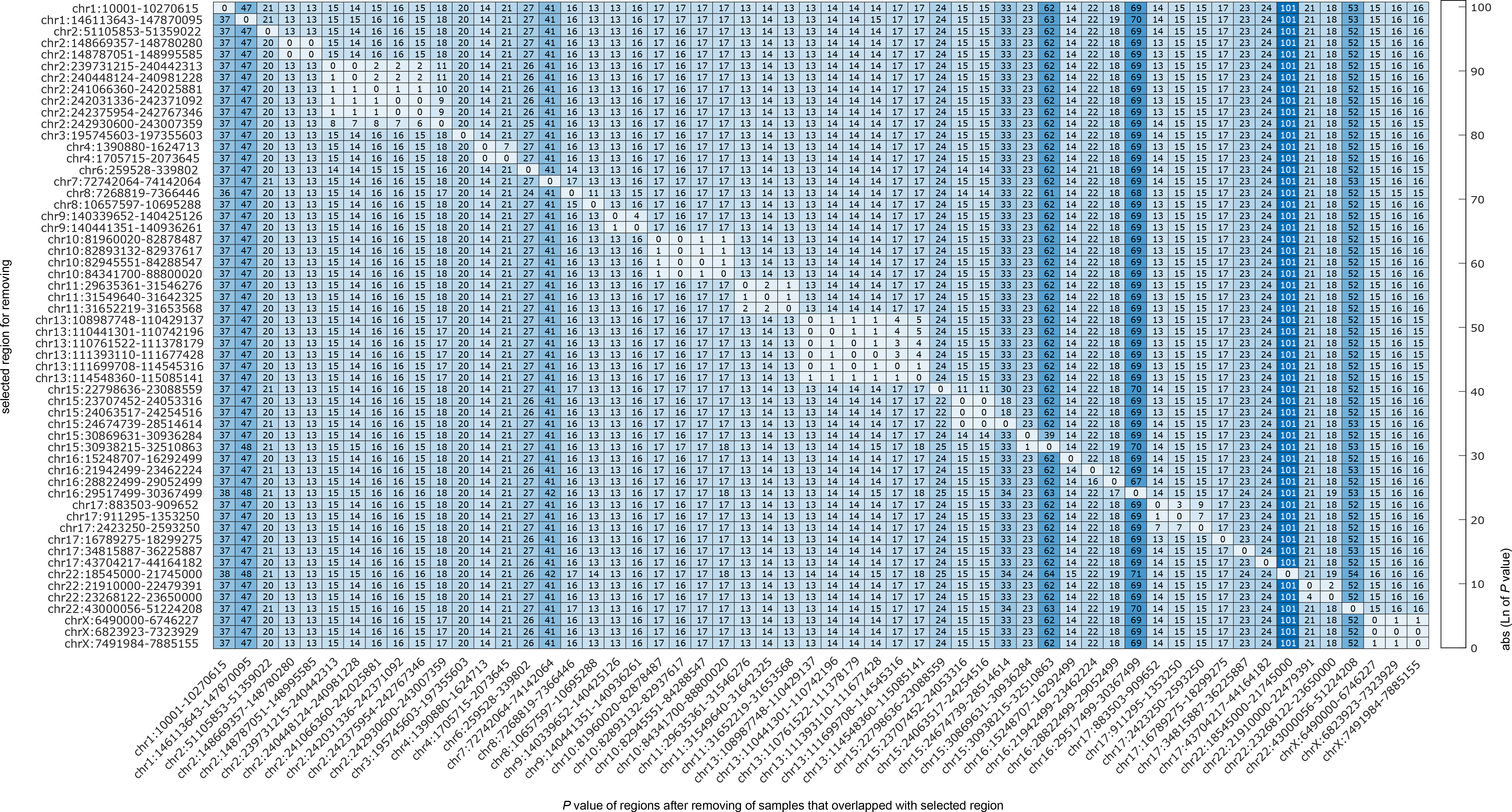
Heatmap matrix of deleted regions after removal. Here, we checked independency/dependency for each region by first removing all case and control samples that overlap the region and then recalculating the *P* value (using one-tailed Fisher exact test) for the remaining 55 regions. This is repeated for all 56 regions. Values in the matrix indicate the natural logarithm of the *P* value after removing all cases and controls that overlap a given region and the y-axis indicates the region selected for removal.

**Figure S7.**
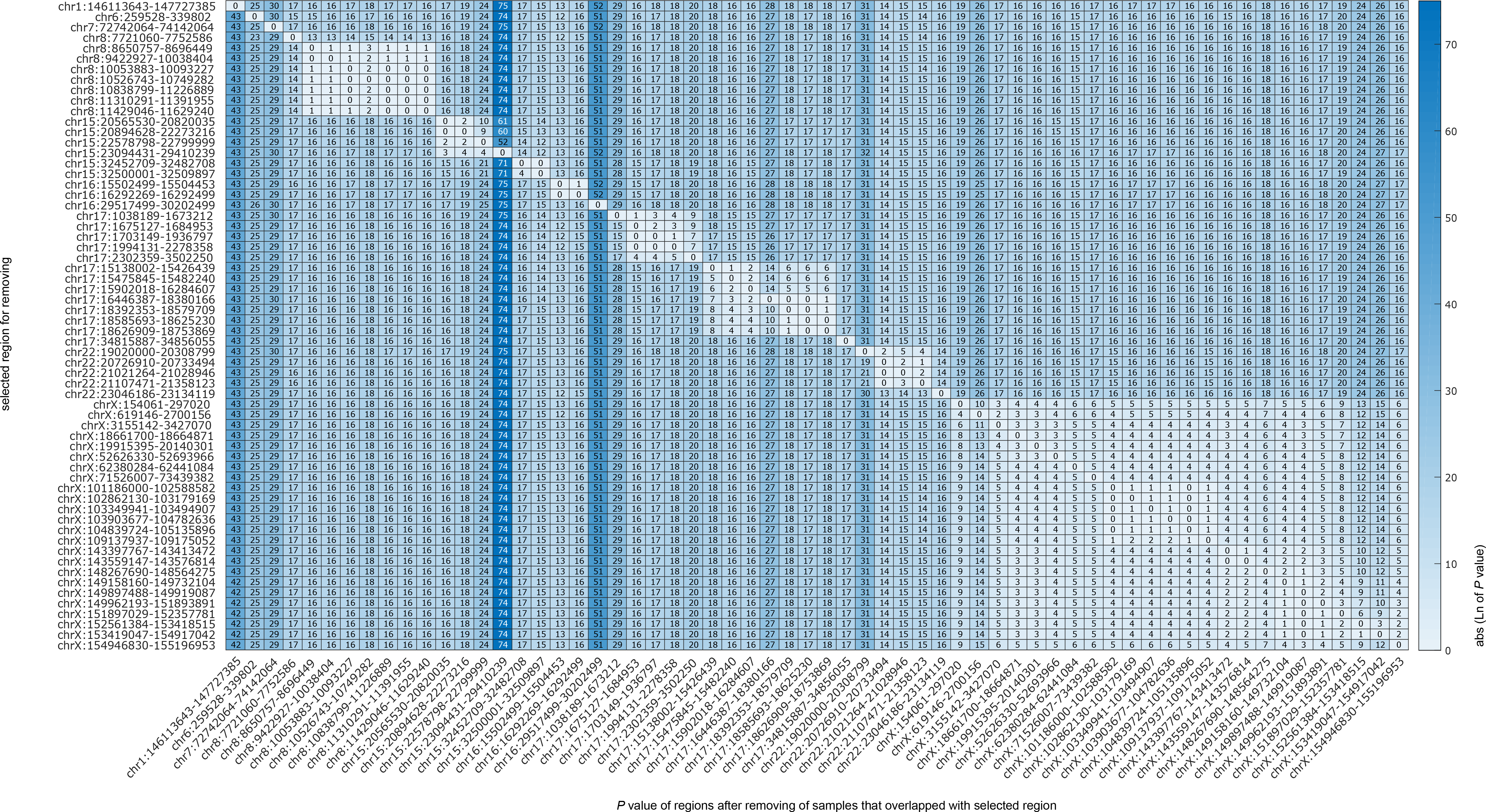
Heatmap matrix of duplicated regions after removal. Here, we checked independency/dependency for each region by first removing all case and control samples that overlap the region and then recalculating the *P* value (using one-tailed Fisher exact test) for the remaining 61 regions. This is repeated for all 62 regions. Values in the matrix indicate the natural logarithm of the *P* value after removing all cases and controls that overlap a given region and the y-axis indicates the region selected for removal.

**Figure S8.**
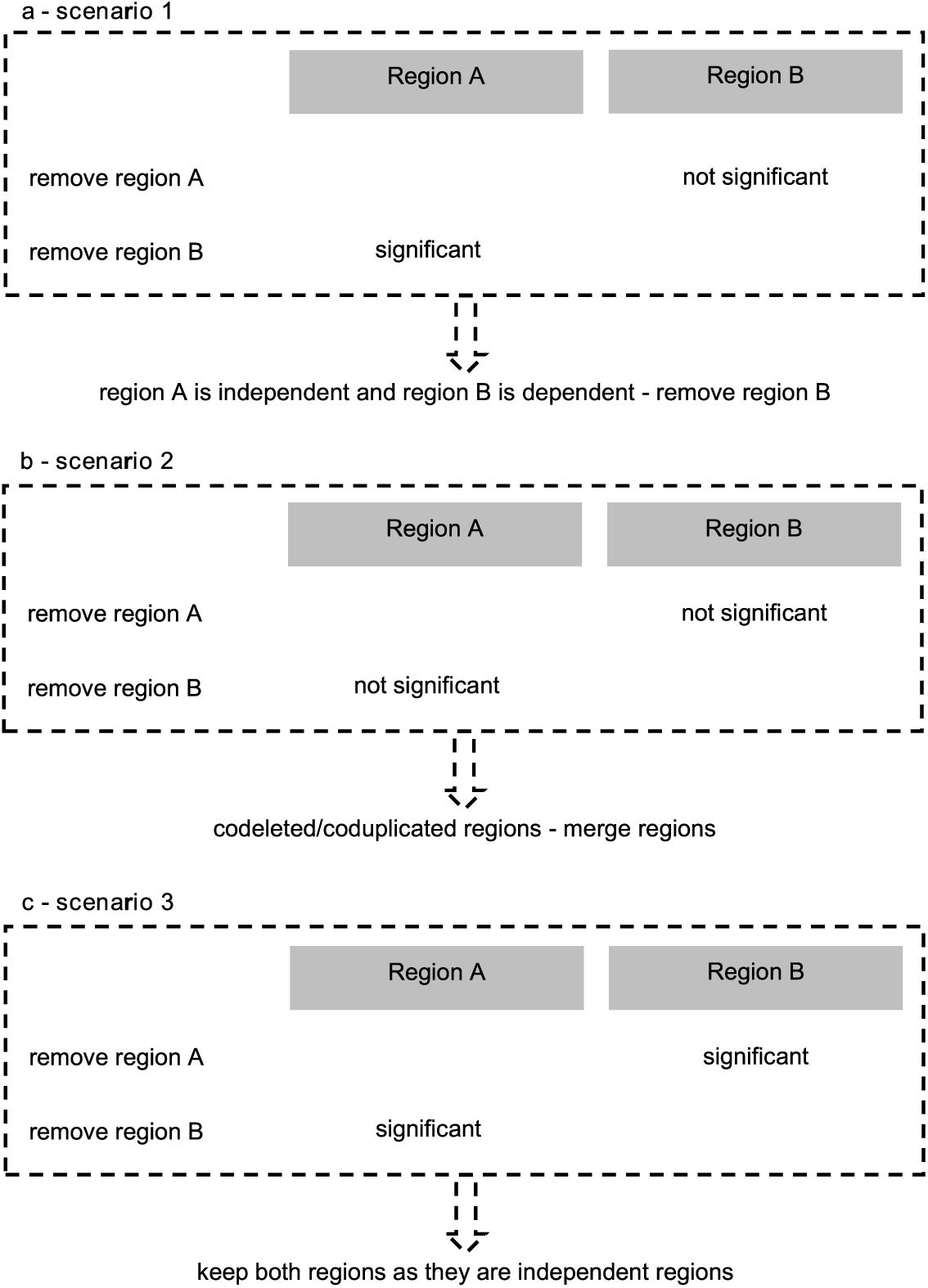
Dependent/independent region scenarios considered. To test independency/dependency of each region we defined three scenarios. **a)** In the first scenario, region A is still significant after removing all case and control samples that overlap with region B, but, region B is not significant after removing all case and control samples that overlap with region A. Here, we keep region A as an independent region and remove region B as a dependent region. **b)** In the second scenario, removal of all case and control samples from region A or region B led to reciprocal lost significance. We call these regions co-dependent regions. For these co-dependent regions, we merged them into a super region. **c)** In the last scenario, removal of case and control samples that overlap with region A or region B still keep another region as a significant region (< 0.002); we refer these regions as independent regions and keep both regions.

**Figure S9.**
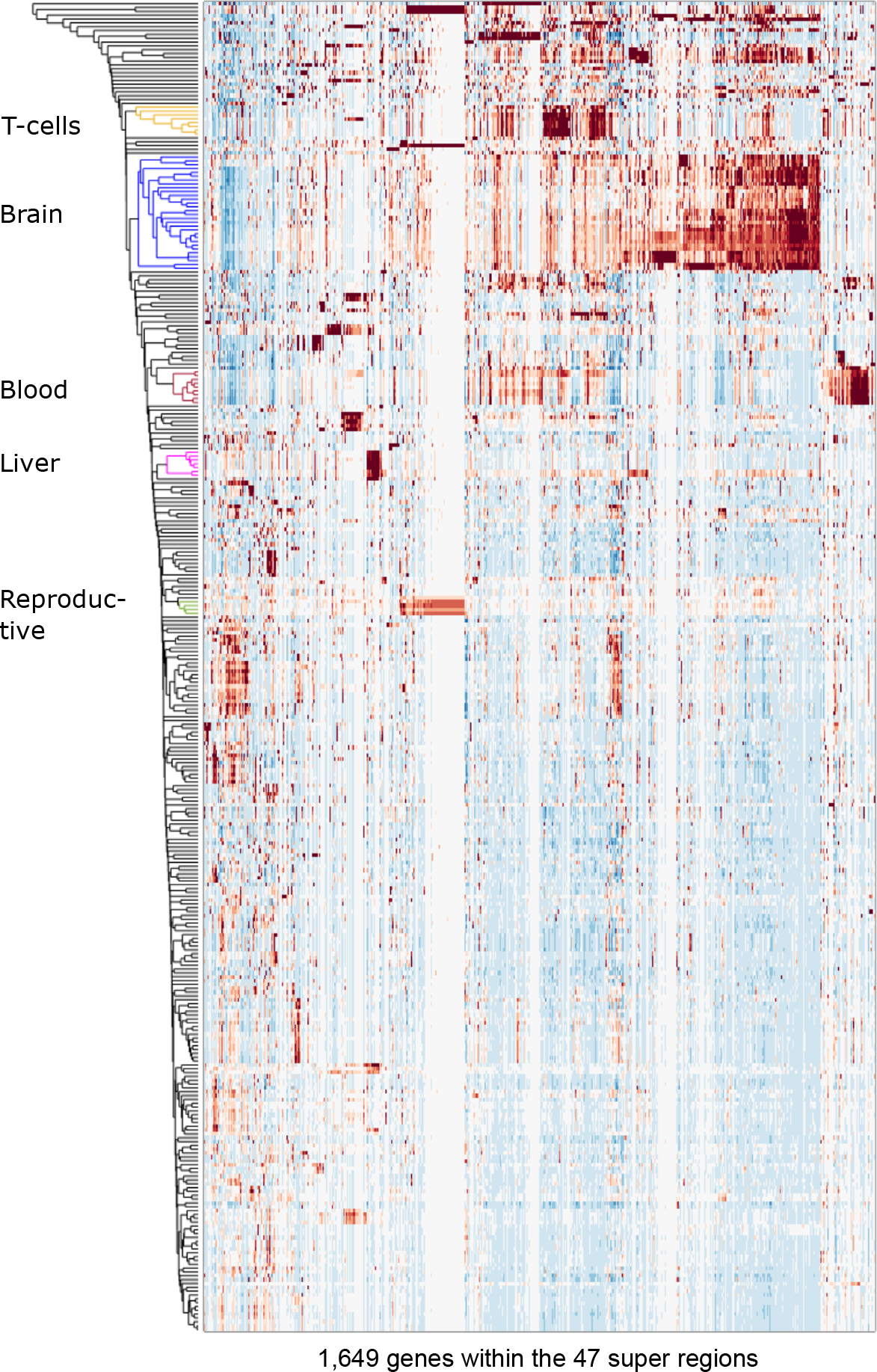
Hierarchical clustering of 1,649 coding and lncRNA genes from the 47 regions based on their average expression across 347 sample ontologies (using the FANTOM CAT expression atlas (19)). Intensity represents the average level of gene expression across all samples annotated with the same sample ontology.

**Figure S10.**
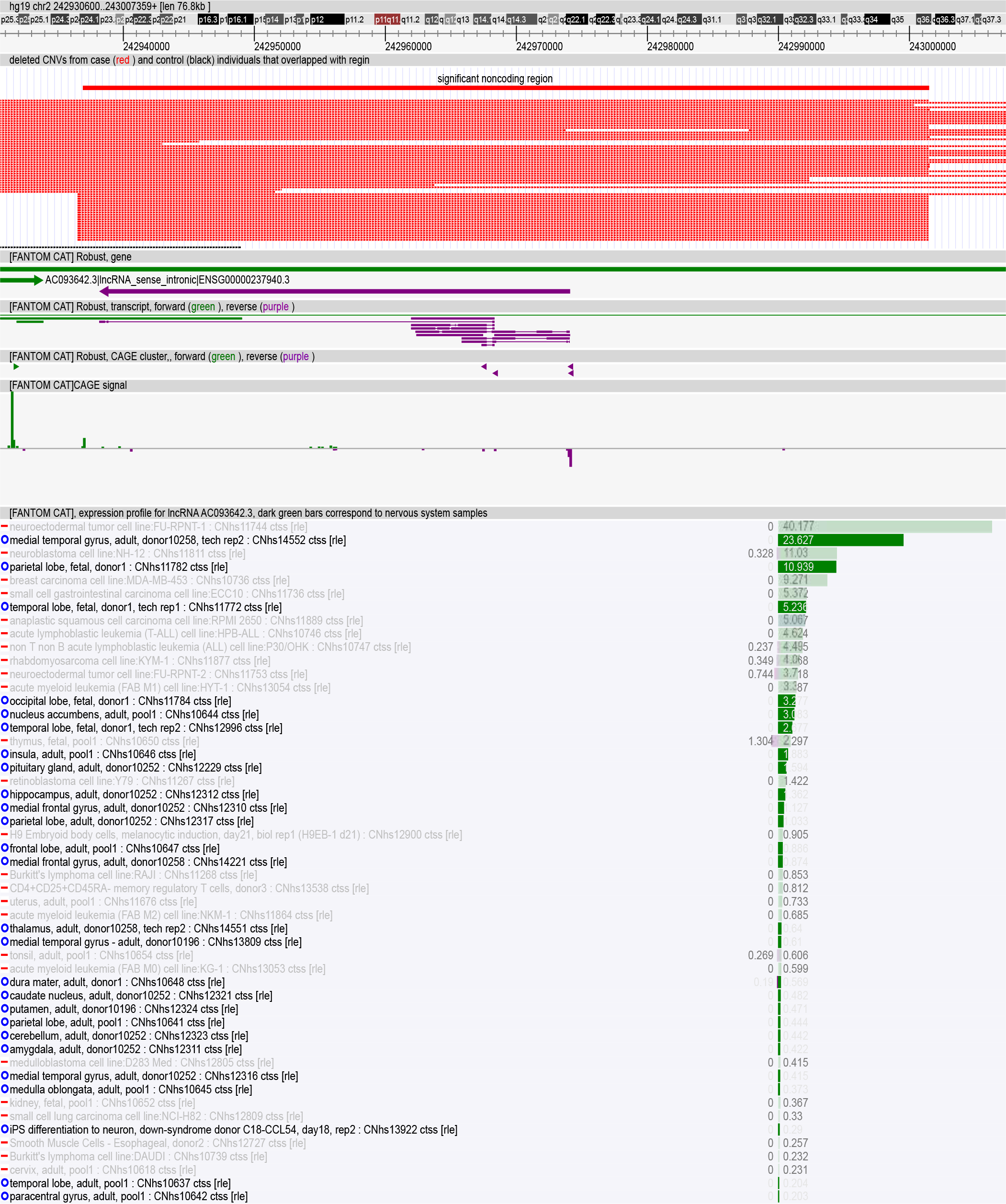
Example of a deleted non-coding region (chr2:242930600-243007359) containing a brain-enriched lncRNA. There is no brain-enriched protein coding gene in this significant regions, indicating the lncRNA is most likely to be responsible against phenotypes associated with this region.

**Figure S11.**
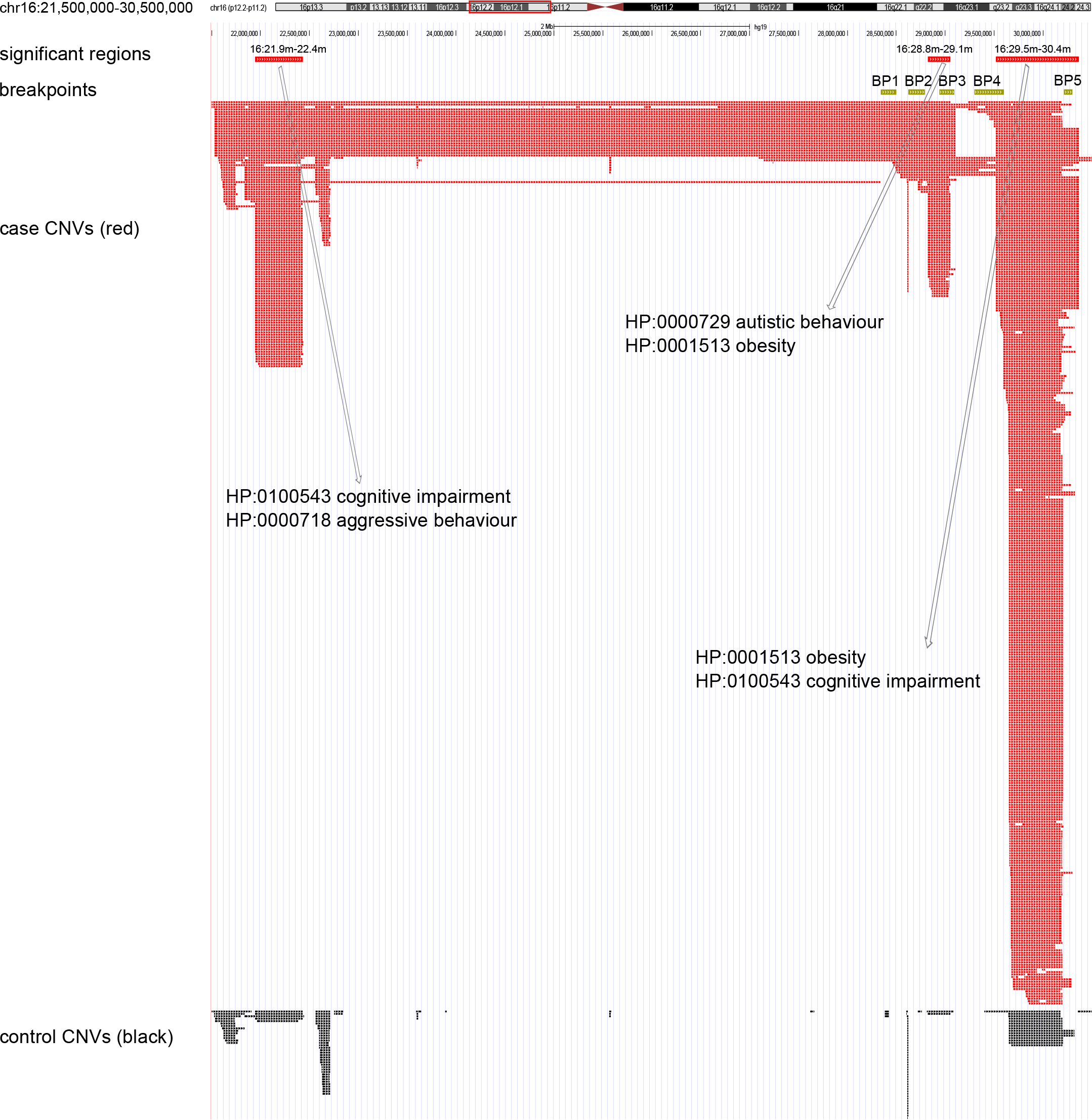
An overview of significant regions (chr16: 21,500,000-30,500,000) identified by SNATCNV and their overlap with breakpoints BP1-BP5. As can be seen the first region chr16:21942499-22462224 is not covered by the regions associated with breakpoints 1 through 5. The second region chr16:28822499-29052499 corresponds to BP2-BP3 and the third region chr16:29517499-30367499 corresponds to BP4-BP5. BP2-BP3 and BP4-BP5 have been previously associated with Autism (44). The image also shows the over-represented human phenotype ontology terms identified from analysis of the phenotypes recorded in DECIPHER.

## Supplementary table legends

**Table S1. List of significant deleted and duplicated regions.** To identify significant regions, we calculated *P* values for 500,000 random permutations of case/control labels to estimate the significant threshold. Here, we reported regions identified at the 99.9999% confidence intervals.

**Table S2. Tools comparison between SNATCNV and existing CNV association tools.** All comparison performed on the same machine with the same number of CPU and the same amount of memory.

**Table S3. Protein coding and noncoding genes in the significant CNV regions identified by SNATCNV (at the confidence interval 99.9999%).** Tissue enrichment *P* value and the best *P* value from permutation analysis provided for each gene in tissues nervous system, nervous system sub-regions, T-cell, muscle cells, mesodermal cell and hematopoietic cell. Induced pluripotent stem cell model of neuron differentiation from FANTOMCAT expression atlas also provided for each gene if any.

**Table S4. Enrichment analysis of genes in the significant regions.** Fraction of genes in recurrent CNV regions with enriched expression in the nervous system, subregions of the brain and non-nervous system samples. We also report enrichment analysis of genes in the significant regions identified by PLINK at its most stringent threshold, in the significant regions identified by Coe *et al.* and in the significant regions identified by Cooper *et al*.

**Table S5. Gene ontology, disease ontology and human phenotype ontology analyses for candidate genes.** Gene ontology, disease ontology, and human phenotype ontology terms with significant enrichment *P* value within candidate genes in both deleted and duplicated regions. We used the gene ontology analysis tool WebGestalt to identify over-representation of ontology terms associated with our genes in the significant regions. We used default values for WebGestalt parameters. Only those ontology terms with more than 5 genes in our regions are shown

**Table S6. Identify independent and dependent regions.** Here, we checked independency/dependency for each region by first removing all case and control samples that overlap the region and then recalculating the *P* value (using one-tailed Fisher exact test) for the remaining 117 regions. For 37 of 118 regions, removal of cases and controls samples found in any other region still had a significant *P* value (< 0.002); we refer to these regions as independent regions. For the remaining 82 regions, 51 lost significance after removal of cases and controls samples from the independent regions; we refer to these regions as dependent regions and removed them from the future analyses. 31 of 118 regions corresponded to co-deleted/co-duplicated scenarios, which means that removal of all cases and controls samples from one region or another led to reciprocal lost significance. For these co-dependent regions, we merged them into 9 super regions. Collectively, after removal process, we identified 47 super regions including 37 independent regions and 9 regions after merging co-dependent regions.

**Table S7. Population analysis for the significant regions.** Here we choose eight top cohorts, which have more than 1000 CNVs in the AutDB database to test if the significant regions are from a specific cohort. We first remove a cohort and then calculate *P* value with one-tailed Fisher exact test for significant regions with the remainder of case and control samples. We repeat this analysis for each of the eight cohort separately.

**Table S8. Literature search for genes within the 47 super regions.** We systematically mined google scholar, SFARI, AutDB, MSSNG and DECIPHER databases to identify known ASD genes. In total, we identified 34 known ASD genes (in 19 different super regions) within significant regions. We also mined the FANTOM-CAT expression atlas and observed that 642 coding genes and 252 lncRNAs in the significant regions are induced in an induced pluripotent stem cell model of neuron differentiation. We also mined the MGI mouse genome informatics database and annotated 186 coding genes that cause nervous system phenotypes when mutated in the mouse.

**Table S9. Literature search for novelty of the 47 super regions.** We systematically mined SFARI, AutDB, MSSNG and DECIPHER databases to annotate 47 super regions as known or novel CNV regions for ASD. In total, 35 regions overlapped with previously identified syndromic/pathogenic regions with ASD related neurodevelopmental phenotypes. For each region, we also provided the number of brain-enriched coding and lncRNA genes, number of brain-enriched coding genes with mice mutant phenotypes that affect the nervous system, and the number of brain-enriched lncRNAs that are induced in an induced pluripotent stem cell model of neuron differentiation.

**Table S10: Sex analysis in the 47 super regions.** Male fold enrichment was calculated by dividing the male fraction by the female fraction for each region. Female fold enrichment calculated through dividing female fraction on male fraction in the region. We note, three of these super regions have been previously reported as a sex bias region.

**Table S11. Phenotypic analysis on the 47 super regions discovered in our study.** To identify significant region-phenotype associations, we only considered those region-phenotype associations that had more than five overlapping patients with a region; we then only counted those associations that a) emerged in at least 5% of patients with an overlapping CNV with the region; b) was 1.5 fold over-represented in samples with CNV overlapped with a super region than DECIPHER patients; c) had a significant fold enrichment more than permutation-specific fold enrichment at 95%C.I. Samples with no phenotype information have been excluded from this analysis.

**TableS12. Brain-enriched genes across the entire genome.** A table of all brain-enriched coding and lncRNA genes identified in the human genome by the one-tailed Fisher exact test and permutation analysis using the FANTOM5 expression data.

**TableS13. List of the 96 known causal protein-coding genes from MSSNG [5] and SFARI [4, 15].** Here, we combined ASD known coding genes from MSSNG consortia with SFARI genes (those genes that had gene score 1 or 2).

